# Determining protein structures using genetics

**DOI:** 10.1101/303875

**Authors:** Jörn M. Schmiedel, Ben Lehner

**Affiliations:** Systems Biology Program, Centre for Genomic Regulation (CRG), The Barcelona Institute of Science and Technology, Dr. Aiguader 88, Barcelona 08003, Spain; Universitat Pompeu Fabra (UPF), Barcelona 08003, Spain; Institució Catalana de Recerca i Estudis Avançats (ICREA), Pg. Lluís Companys 23, 08010 Barcelona, Spain

## Abstract

Determining the three dimensional structures of macromolecules is a major goal of biological research because of the close relationship between structure and function. Structure determination usually relies on physical techniques including x-ray crystallography, NMR spectroscopy and cryo-electron microscopy. Here we present a method that allows the high-resolution three-dimensional structure of a biological macromolecule to be determined only from measurements of the activity of mutant variants of the molecule. This genetic approach to structure determination relies on the quantification of genetic interactions (epistasis) between mutations and the discrimination of direct from indirect interactions. This provides a new experimental strategy for structure determination, with the potential to reveal functional and *in vivo* structural conformations at low cost and high throughput.

## Introduction

Mutations within a protein or RNA can have non-independent effects on fitness ^1-4^. Indeed, the effects of double mutants have long been used to probe the energetic couplings between positions in a protein to understand determinants of protein folding and stability ^5,6^. Early work revealed that at least some strongly interacting positions within a protein are in direct structural contact ^5-8^. Deep mutagenesis of proteins ^9-12^ and RNAs ^13-16^ has further confirmed this conclusion that some – but by no means all – genetic (or epistatic) interactions occur between structurally proximal mutations.

Further support for the idea that non-independence between mutations provides structural information comes from the analysis of amino acid and nucleotide sequence evolution. Here, correlated pairs of amino acids or nucleotides in multiple sequence alignments identify co-evolving positions within proteins and RNAs ^17-19^. These patterns of co-evolution have been used to identify energetically-coupled positions and ‘sectors’ within proteins ^20,21^. Moreover, when very large numbers of homologous proteins and RNAs are avaiable in sequence databases, the application of global statistical models has proven sufficient to discriminate direct structural contacts from patterns of co-evolution ^22-24^, allowing the prediction of macromolecular structures and interactions ^25-34^.

Could epistatic interactions quantified from deep mutational scanning experiments be used to determine macromolecular structures? If successful, structure determination by deep mutagenesis would offer a number of advantages over established techniques. First, it requires no specialized equipment or expertise beyond the ability to mutate a molecule, select functional variants, and quantify enrichments by sequencing. Appropriate *in vitro* and *in vivo* selection assays already exist for very many molecules of interest and generic assays based on folding, stability, and physical interactions have also been developed ^9,35-38^. Second, it could be applied to molecules whose structures are difficult to determine by physical techniques such as intrinsically disordered and membrane proteins. Third, unlike evolutionary coupling analysis there is no requirement for large numbers of homologous sequences and so it could be applied to fast-evolving, recently-evolved and *de novo* designed proteins and RNAs ^26,32,39^. Finally, and perhaps most importantly, it would provide a general strategy to determine the physiologically relevant structures of molecules whilst they are performing particular functions that can be selected for, including *in vivo* within cells. A cheap and straightforward approach for studying macromolecular structures *in vivo* would be a very exciting new frontier for cell biology.

Here we show that deep mutational scanning (DMS) of proteins can provide sufficient information to determine their high-resolution three-dimensional structures. Our statistical approach quantifies how often mutations between positions interact epistatically and how such epistatic interaction patterns correlate. These metrics accurately identify individual tertiary structure contacts as well as secondary structure elements within a protein. The same approach also identifies contacts between protein interaction partners. DMS data alone suffices to determine protein structures with accuracies down to 1.9Å backbone root mean square deviation (RMSD) compared to known reference structures. Moreover, we show that deep learning can further improve prediction performance, allowing the use of much sparser and lower quality DMS datasets for structure determination. This approach therefore provides a new experimental strategy for structure determination that can reveal functional and *in vivo* structural conformations at low cost and high throughput.

## Results

### Epistasis is enriched in but not exclusive to structural contacts

To investigate how genetic – or epistatic – interactions between mutations in a protein relate to structure we first used deep mutational scanning data for the immunoglobulin-binding protein G B1 domain (GB1) generated by Olson, et al. ^11^. This dataset is the most complete double mutant deep mutagenesis of a protein domain reported to date and was generated by replacing each of 55 residues of the wild-type domain with all 19 alternative amino acids both individually and as double mutant pairwise combinations, resulting in a library of more than half a million variants (55*19 = 1,045 single mutants plus nearly 55*54/2*19*19 = 536,085 double mutants). mRNA display was used to combine an *in vitro* immunoglobulin G binding assay with a sequencing readout to determine protein fitness via changes in variant frequencies in the library before and after binding (Extended Data Figure 1, steps 1-4); resulting in a two orders of magnitude measurement range with a median relative error of fitness estimates of 2.8% (see Figure 2A, Table 1 and Methods).

**Table 1:** Dataset properties

| Dataset | Mutated positions | % double mutants<sup>\$</sup> | % doubles quantifiable<sup>#</sup> | | # input reads per double mutant (median)<sup>*</sup> | measurement range (log fitness units)<sup>+</sup> | relative error (median)<sup>&</sup> |
| --- | --- | --- | --- | --- | --- | --- | --- |
|  |  |  | positive epistasis | negative epistasis |  |  |  |
| Protein G B1 domain | 55 | 97 | 80 | 55 | 248 | 5.8 | 2.8% |
| hYAP WW domain | 33 | 10 | 8.3 | 0.8 | 73 | 0.8 | 8.6% |
| Pab1 RRM2 domain | 25 | 11 | 8.3 | 3.9 | 209 | 3.1 | 3.7% |
| FOS-JUN | 2 x 32 | 43 | 37 | 31 | 124 | 8.6 | 3.6% |

<sup>\$</sup> median percentage of all possible double mutants (361 per position pair) that passed read quality thresholds per position pair
<sup>#</sup> median percentage of all possible double mutants (361 per position pair) that passed read quality thresholds and are deemed suitable for epistasis quantification per position pair
<sup>\*</sup> summed number of reads across all input replicates for double mutants that passed read quality thresholds
<sup>+</sup> measurement range of selection assay: log fitness range between peak of lethal mutants and the wild-type variant
<sup>&</sup> median error of fitness estimates of double mutant variants relative to measurement range of selection assay

We first computed which double mutant variants show epistatic fitness effects, i.e. non-independent fitness effects of the constituting single mutant variants (Figure 1B). Non-specific dependencies between mutants might be introduced by non-linearities in the fitness assay, systematic biases in error magnitudes as well as non-specific epistatic behavior, e.g. from thermodynamic stability effects ^1,9^. We thus applied a non-parametric null model - the running median of double mutant fitness values given the constituting single mutant fitness values - for the independence of mutations. Equivalently, we calculated 5th and 95th percentile fitness surfaces; and classified double mutants with fitness lower than the 5^th^ percentile as negative epistatic and double mutants with fitness higher than the 95^th^ percentile as positive epistatic. We restricted the evaluation of positive or negative epistasis, however, to specific subsets of the data, where measurement errors do not impede epistasis classification (Extended Data Figure 2C, see Methods), which results in about 80% and 55% of double mutants being suitable for positive or negative epistasis classification, respectively, with a lot of variability across the position matrix (Extended Data Figures 2D-F and Table 1).

**Figure 1:**
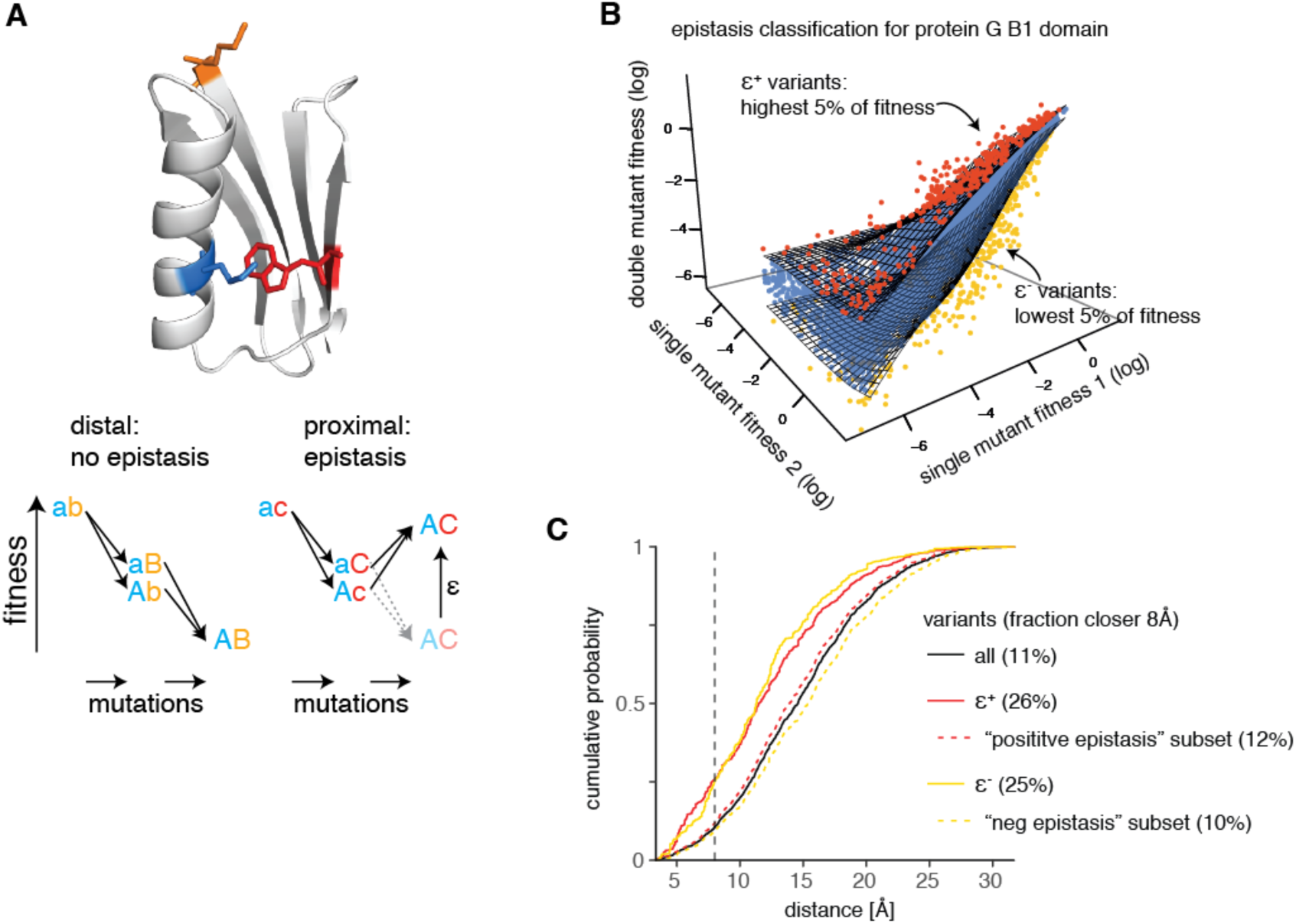
Extracting epistatic mutational effects from deep mutational scanning of a protein domain. A. Premise: If epistatic interactions relate to structural contacts then quantifying epistatic interactions should suffice to predict a molecule’s structure. Structure: protein G B1 domain (PDB entry: 1pga) with residues a, b, and c colored. B. Classifying epistatic variants based on deviations from expected fitness (based on quantile fitness surface approach). Variants above the 95th or below the 5th percentile of double mutant fitness given their single mutant fitness values were classified as positive (red, ε^+^) or negative (yellow, ε^−^) epistatic, respectively. Shown is a random sample of 10^4^ variants in GB1 domain ^11^. C. Distance distribution of epistatic variants separated by more than 5 amino acids in the linear sequence. (side-chain heavy atom distance in reference structure). Positive and negative epistasis subsets refer to the sets of variants applicable for epistasis analysis (see Extended Data Figure 2C). All variants, n = 400647; positive epistatic variants ε^+^, n = 14127; positive epistasis subset, n = 315862; negative epistatic variants ε^−^, n = 9837; negative epistasis subset, n = 208442.

Consistent with previous observations ^10-12^, both positive and negative epistatic double mutants are enriched for proximal variants, for example, more than 2-fold at 8Å distance (side-chain heavy atom minimal distance, Figure 1C). However, about 75% of epistatic interactions are between positions that are not in direct contact in the tertiary protein structure (as judged by an 8Å distance cutoff), suggesting that indirect effects often underlie epistatic interactions within a molecule ^20,21^. The challenge for structure determination therefore becomes how to infer direct structural contacts from the mixture of direct and indirect effects that must underlie epistasis.

### Aggregated epistatic interactions predict tertiary structure contacts

To distill direct contacts from a list of thousands of epistatic double mutants we first aggregated epistatic information on the amino acid position-pair level by calculating the fraction of positive or negative epistatic double mutant variants per position pair (Figure 2A).

**Figure 2:**
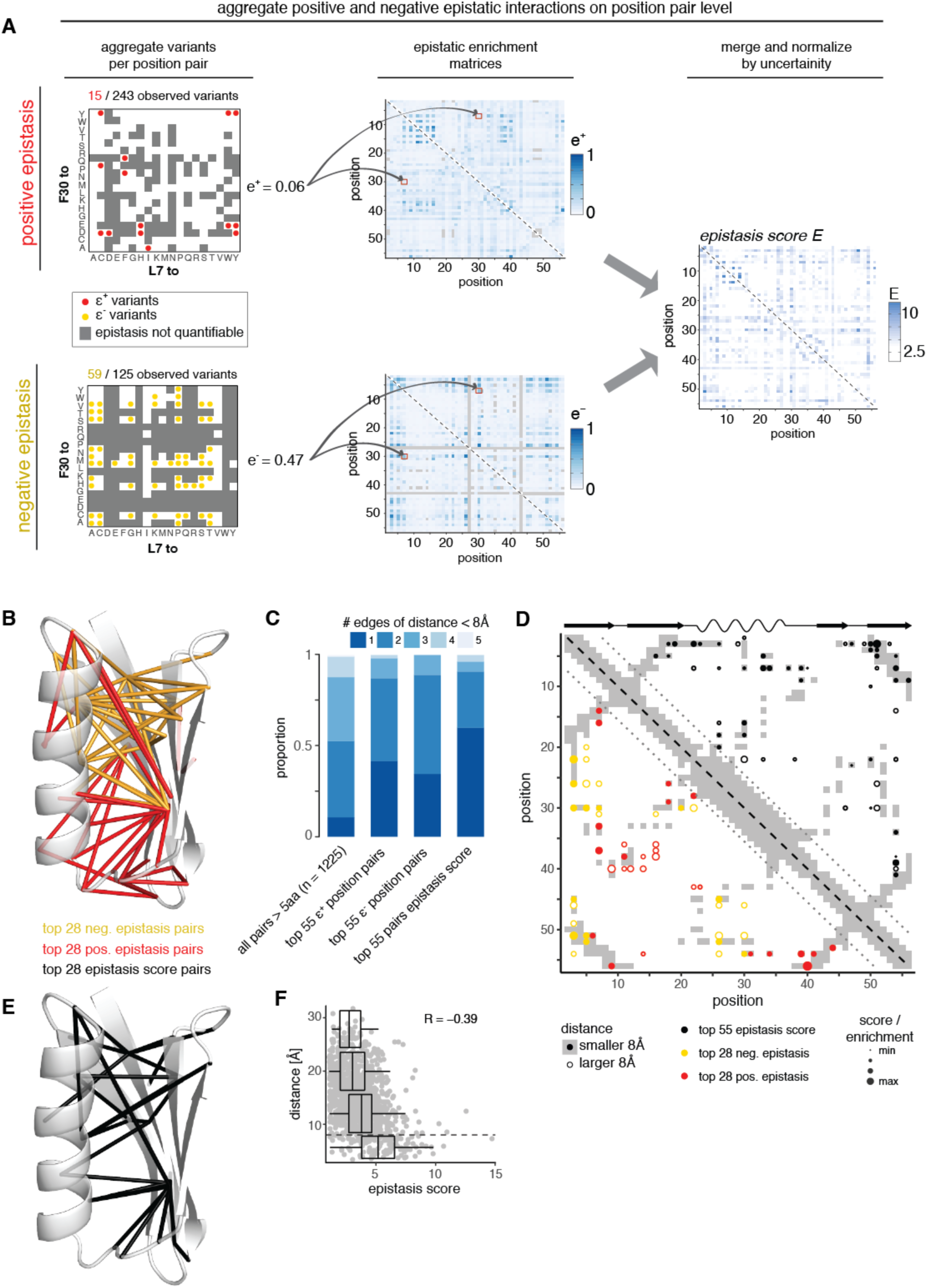
Aggregated epistasis scores enrich for direct structural contacts. A. Workflow for aggregating positive and negative epistatic interactions on the position-pair level and merging them into a final *epistasis score*. B. Top 28 position pairs (> 5 amino acids in linear sequence) each with highest positive (red) and negative (yellow) epistatic fractions marked on the reference structure (PDB entry 1pga). C. Minimal number of edges (contact with distance < 8Å in reference structure) connecting position pairs. One edge – positions are direct contacts, two edges – positions have a common contact and so forth. D. Interaction score map for top 28 position pairs each with highest positive (red) and negative (yellow) epistatic fractions (lower left triangle) and top 55 position pairs with highest *epistasis score* (upper right triangle). Dot size indicates relative epistatic enrichments or score; dot fill indicates distance below 8Å. Underlying in grey is the contact map of the reference structure (PDB entry 1pga, distance < 8Å) and shown on top its secondary structure elements (wave – alpha helix, arrow – beta strand). E. Top 28 position pairs (> 5 amino acids in linear sequence) with highest *epistasis scores* marked on the crystal structure (PDB entry 1pga). F. Distance of position pairs as a function of *epistasis scores*. Boxplots are spaced in distance intervals [0,8), [8,16), [16,24) and [24,32) Å. Dashed horizontal line indicates 8Å. Pearson correlation coefficient is indicated.

In the GB1 epistasis dataset even moderate enrichments for positive and negative epistatic variants are mutually exclusive (Extended Data Figure 3A). Moreover, the strongest positive and negative epistatic enrichments are separated in two clusters of proximal positions in the protein that exhibit mostly either positive or negative interactions among themselves, but hardly any epistatic interactions between clusters (Figures 2B,D and Extended Data Figure 3B), as also noted before by Olson, et al. ^11^.

Consistent with epistatic interaction clusters forming a dense network of proximal positions, we find that, of the top 55 epistatic pairs, 42% and 35% are direct contacts (connected by one edge smaller than 8 Ångström (Å), 3.9 and 3.2-fold over expectation) and another 45% and 55% share a common neighbor (connected via two edges < 8Å), for positive and negative epistatic interactions respectively, while interactions across more edges are depleted (Figure 2C; throughout the manuscript we only consider position pairs spaced by more than 5 amino acids in the linear sequence; closer positions are trivially also close in 3D space, and their proximity contributes little to successful structure prediction ^28^).

While aggregation of epistatic information between position pairs thus better discriminates structural contacts than individual epistatic interactions, positive and negative epistatic interactions still contain disparate structural information of the protein domain. We therefore merged positive and negative epistatic information by computing the weighted averages of epistatic fractions per position pair given their uncertainty due to fitness measurement errors and the finite number of observed double mutant variants via a resampling approach (Extended Data Figure 1 and Methods). A final *epistasis score* per position pair was obtained by normalizing these weighted averages by their uncertainty (a z-score), thus giving priority to position pairs with high confidence enrichments.

The position pairs with highest *epistasis scores* are well distributed across the domain (Figures 2D,E), and the number of direct contacts (one edge < 8Å) among the top 55 *epistasis score* pairs increases to 60% (Figure 2D), thus showing that the *epistasis score* successfully incorporates information from both positive and negative epistasis to discriminate direct contacts. Moreover, direct contacts as a whole are enriched for high *epistasis scores*, while further away position pairs show a gradual decrease of *epistasis scores* (Pearson correlation coefficient R = -0.39, p < 10^-6^, Figure 2F).

Thus, although many interactions are indirect, physical contacts are an important determinant of epistasis and aggregating information on position pairs and merging positive and negative epistasis information better discriminates these direct structural contacts across the protein domain.

### Tertiary structure neighborhood leads to correlated epistatic patterns

If epistasis arises mainly from structural interactions, a position’s epistatic interaction profile with all other positions in the protein should provide a signature of its structural location (Figure 3A). Comparing these signatures between positions should thus reveal structurally close positions - similar to how correlated epistasis profiles in genetic interaction networks serve to identify physical and functional interaction partners ^40^.

**Figure 3:**
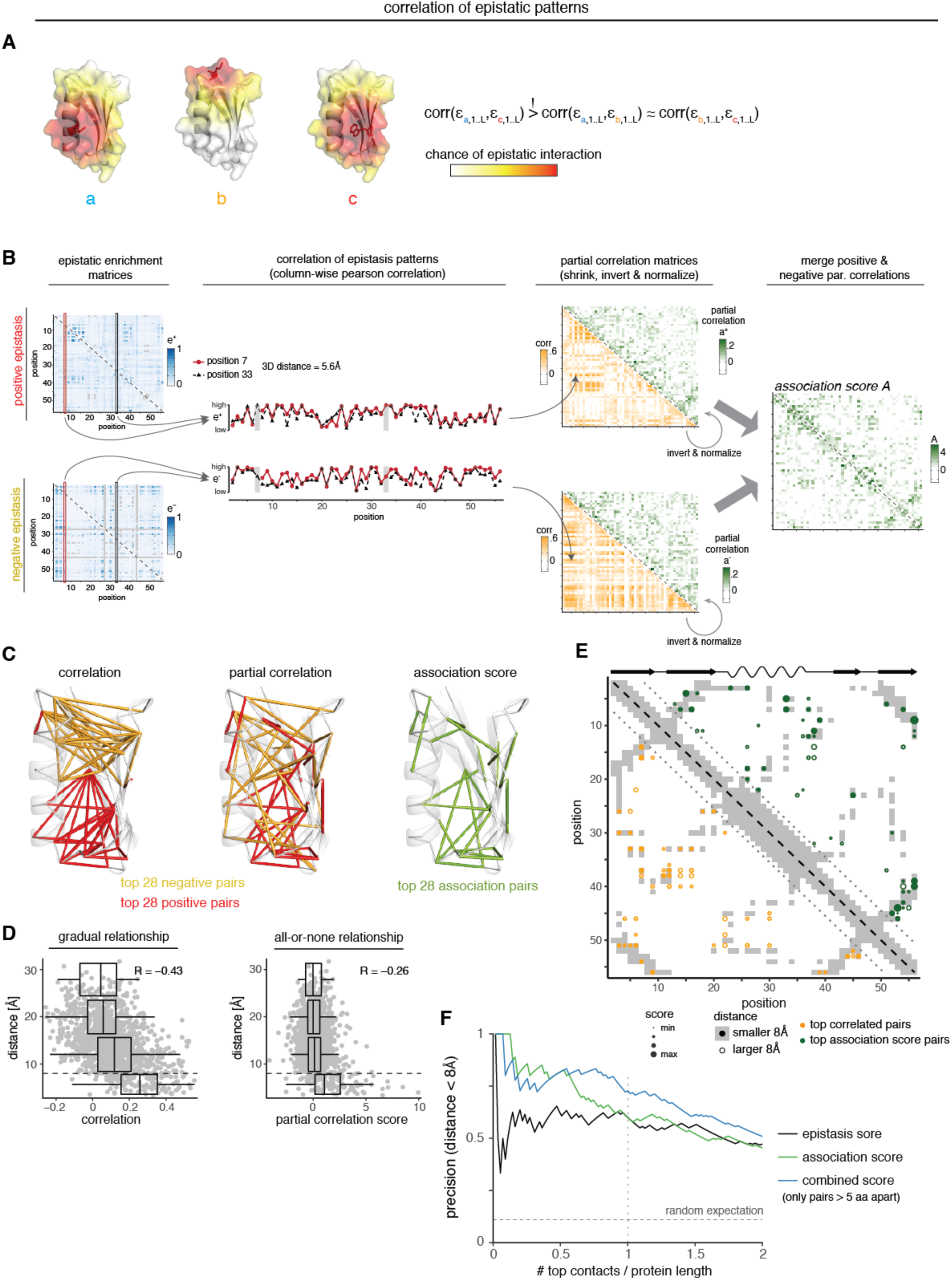
Tertiary structure neighborhood leads to correlated epistatic patterns. A. Mutations in directly contacting residues should interact similarly with all other mutations in the protein. Thus, the similarity of epistasis patterns of two positions with all other positions in the protein should inform about their structural proximity. B. Workflow for quantification of correlated epistasis patterns. Pairs of columns from epistatic enrichment matrices (here columns 7 and 33) are compared and their Pearson correlation coefficients are calculated, which constitute entries in the correlation matrix (here entries 7:33 and 33:7, due to matrix symmetry). Correlation matrices are inverted to yield the partial correlation matrices. Finally, entries of the positive and negative partial correlation matrices are merged (weighted average by uncertainty) and z-normalized to yield *association scores* (see Methods). C. Top 28 position pairs (> 5 amino acids in linear sequence) marked on reference structure. Left: Top pairs from positive (red) or negative (yellow) epistasis pattern correlations. Middle: Top pairs after partial correlation transformation. Right: Top *association score* pairs (merge positive and negative partial correlations). D. Distance of position pairs as a function of merged correlation (left) or *association scores*. Boxplots are spaced in intervals of 8Å. Dashed horizontal line indicates 8Å. Pearson correlation coefficient is indicated. E. Interaction score map for top 55 position pairs with highest merge correlation (positive and negative correlations merged, lower left triangle, orange) and *association scores* (upper right triangle, green). Dot size indicates relative correlations or scores; dot fill indicates distance below 8Å. Underlying in grey is the contact map of the reference structure (PDB entry 1pga, distance < 8Å) and shown on top its secondary structure elements (wave – alpha helix, arrow – beta strand). F. Precision of interaction scores to predict direct contacts (distance < 8Å in crystal structure 1pga) as a function of top scoring position pairs. Only position pairs with linear chain distance greater than 5 amino acids are considered (n = 1225 pairs, n = 131 direct contacts in reference structure). Horizontal dashed line indicates random expectation.

To test the idea that pattern correlation should reveal structural proximity, we calculated the correlations between the epistatic enrichment vectors for all position pairs (Figure 3B). Consistently, pair-distances and similarity of epistasis patterns between positions are strongly correlated (Pearson correlation coefficient = -0.43, p < 10^-6^, n = 1225, Figure 3D). Top correlated pairs from positive or negative interaction patterns do, however, form mutually exclusive clusters within the protein domain that are nearly identical to the clusters observed for direct positive and negative interactions (Figure 3C and E, c.f. Figure 2B). Thus, while correlations of epistatic interaction patterns are a good indicator of distance within the protein structure, they suffer from the same issues as epistatic enrichments, namely poor discrimination of direct and indirect interactions and disparate structural information.

We reasoned that partial correlations - the association between two positions after accounting for the global correlation structure - might provide the possibility to eliminate the dependencies observed in the epistasis pattern structure and thus help to distinguish direct from indirect contacts; similar to how mean-field approaches can help discriminate direct from indirect evolutionary couplings in multiple sequence alignments ^22,28,41^. We derived partial correlations by inversion of the correlation matrices, merged values from positive and negative epistatic patterns by their estimated uncertainty, and ranked these merged values by their z-scores, which we refer to as *association scores* (Figure 3B and Methods).

In contrast to the correlation of epistasis patterns, partial correlation of epistasis patterns for both positive and negative epistasis display no clustering but are well distributed across the whole protein domain, consistent with partial correlations removing dependencies between correlated pairs (Figure 3C). Moreover, the merged *association scores* are less well correlated with pair-distance (Pearson correlation coefficient R = -0.26, p < 10^-6^, n = 1225) and show a more binary all-or-none response, with most distant position pairs having an *association score* around 0 and only proximal pairs systematically deviating to higher values (Figure 3D). Moreover, the top pairs involve many different individual positions and are well distributed across the protein domain (Figure 3C and E). Thus, *association scores* are able to prioritize direct over indirect structural contacts across the whole protein domain.

### Combining *epistasis* and *association* scores better discriminates structural contacts

We derived a *combined score* by summing the standardized *epistasis* and *association scores*, to explore whether combining information from individual epistatic interactions and epistasis interaction patterns can improve proximity estimates; thereby prioritizing position pairs that are both enriched for direct epistatic interactions and have correlated epistasis patterns.

We evaluated the precision of the three interaction scores in predicting direct contacts in the protein domain. For all pairs separated by more than 5 amino acids in the linear sequence, the *epistasis score* has a roughly constant precision of around 60% across the first 2*L predicted contacts (L being the mutated length of the protein i.e. 55 amino acids). The *association score* has higher precision than the *epistasis score* up to the first L predicted contacts, with a precision of 79% at L/2 top contacts. Finally, the *combined score* has similar precision to the *association score* for the first L/2 contacts, but then remains at higher precision, with an improvement of about 10-15% over the individual scores at more predicted contacts (73% at L contacts); showing that combining information from epistatic interactions and interaction patterns further improves the discrimination of direct structural contacts.

Together, the derivation of the interaction scores demonstrates that it is possible to discriminate direct three-dimensional structural contacts from a mainly non-proximal set of epistatic interactions within a protein domain.

### Periodic epistatic patterns reveal secondary structure arrangements

We investigated whether the periodic geometrical arrangement of amino acid residues in secondary structures results in periodic epistasis patterns ^26,42^. Within an alpha helix with 3.6 residues per helical turn, a helical position would be predicted to interact epistatically with the third or fourth-over position along the linear amino acid chain (Figure 4A). Equivalently, within a beta strand, positions should interact epistatically with the next-but-one position.

**Figure 4:**
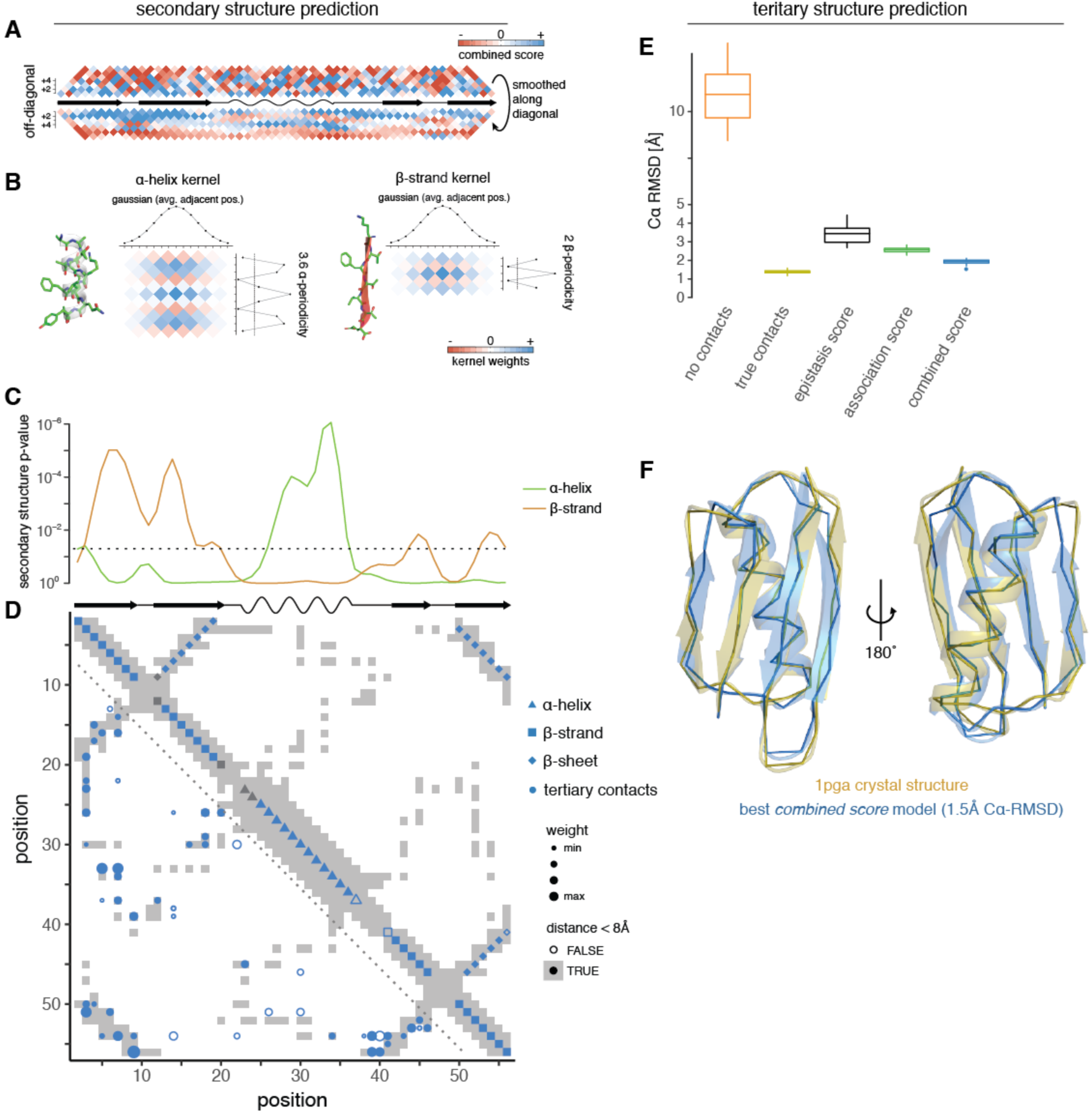
Secondary and tertiary structure prediction from deep mutational scanning data. A. Local interactions reveal signatures of secondary structure elements. Middle line is diagonal of interaction score map (rotated by 45 degree) and shows secondary structure elements of reference structure (PDB entry 1pga). Data above diagonal shows *combined score* data close to the diagonal, i.e. local interactions. Below the diagonal, the same data are smoothed with a Gaussian kernel along the direction of the diagonal (i.e. horizontally, length of Gaussian kernel as for kernels in panel b) to reveal periodicities in local interactions. B. Two-dimensional kernels for alpha helix and beta strand detection. Kernel has a sinusoidal or alternating profile in the off-diagonal direction to detect alpha helices and beta strands propensities, respectively and a Gaussian profile along the diagonal, to average over propensities of adjacent positions. C. Secondary structure propensity derived from kernel smoothing (orange – beta strand, green – alpha helix). P-values were derived by comparison to randomized datasets (see Methods). Dashed line indicates p = 0.05. D. All structural predictions derived from *combined score* data. Lower left: Top 55 non-local (>5 aa in linear sequence) position pairs, i.e. tertiary contacts (circles); fill indicates correct prediction at 8Å, size of circles indicates relative score. Upper right: Predicted secondary structure elements (triangle – alpha helix, square – beta strand, diamond – beta sheet interaction). Fill indicates correct prediction. Note that beta strand predictions are derived by intersection of beta strand propensity (as shown in panel C) and results from beta sheet prediction (Extended Data Figure 4B, see Methods). Underlying in grey is the contact map of the reference structure (PDB entry 1pga, distance < 8Å) and shown on top are its secondary structure elements (wave – alpha helix, arrow – beta strand). E. Accuracy (*C*α root-mean-square deviation) of top 5% structural models generated from deep mutational scanning data derived restraints compared to GB1 reference structure. Structural models were generated in XPLOR-NIH by simulated annealing with restraints derived from top 55 top scoring position pairs, secondary structure element prediction and beta sheet pairing predictions from the indicated interaction scores. No contacts – negative control with restraints only for secondary structure (predicted by PSIPRED)^43^. True contacts – positive control with 55 contacts (random subset), secondary structure elements and beta sheet interactions restraints derived from reference structure. F. Overlay of top structural model generated with restraints from *combined score* (blue) and crystal structure (gold, PDB entry 1pga). Shown is backbone ribbon and secondary structure cartoon generated in PyMOL ^44^.

We used a two-dimensional kernel smoothing approach to estimate the positions of alpha helices and beta strands from the deep mutational scanning data (Figure 4B). Here, the propensity of a position to belong to an alpha helix or a beta strand depends on whether it shows the expected periodicity in its interaction with neighboring positions, as well as whether neighboring positions display similar propensities for the same secondary structure element, and how strong these interactions are compared to those found in randomized data sets (see Methods).

We found that, while secondary structure element predictions derived from direct interaction- based *epistasis scores* are somewhat inaccurate and underpowered, predictions derived from correlation-based *association scores* (as well as *combined scores*) coincide very well with secondary structure elements in the reference structure (Figure 4C and Extended Data Figure 4C), with precision and recall values of about 90% (Extended Data Figure 4D). This suggests that the correlated profiles of epistatic interactions are informative about side chain orientations and also that eliminating transitive interactions is important for (secondary) structure prediction.

We further used two-dimensional kernel smoothing to detect parallel and anti-parallel beta sheet interactions, by applying beta strand kernels to off-diagonal entries on the interaction score matrices (Extended Data Figure 4A, see Methods). Several stretches of position pairs show the expected alternating interaction profiles for either parallel or anti-parallel beta sheets (Extended Data Figure 4B), with the top predictions corresponding to the known beta-sheet interactions in the reference structure (Figure 4D). Furthermore, updating beta strand predictions according to inferred beta sheet pairings can further improved beta strand prediction itself, notably introducing a correct split of beta strand 1 and 2 and adjusting the length of beta strands 3 and 4 (Figure 4C,D).

Together this shows that epistatic interaction data contains information on the periodic secondary structure of a protein domain and, *vice versa*, that secondary structure strongly influences genetic interactions.

### Protein structure determination by deep mutagenesis

Together, these findings show that deep mutagenesis data contain substantial information about a protein’s secondary and tertiary structure. We therefore tested whether the data would suffice to determine *ab initio* the structure of protein G domain B1. We performed structural simulations by simulated annealing using the XPLOR-NIH modeling suite ^45^, with structural restraints derived from the deep mutational scanning data (see Methods). In particular, we defined distance restraints (distance < 8Å between Cβ atoms) for the top scoring position pairs; we found that using the top L (L = 55) contacts gave best results (Extended Data Figure 4F). Furthermore, we defined dihedral angle restraints for predicted secondary structure elements. Finally, we defined restrictive distance restraints (distance smaller than 2.1Å for N-H : C=O atom pairs) for beta sheet positions that form hydrogen bonds with each other.

We evaluated the top 5% of structural models (25/500, evaluation based on XPLOR internal energy terms) generated against the known crystal structure of protein G domain B1 (PDB entry 1pga) (Figures 4E and Extended Data Figure 4F). Models predicted from *combined score* data performed best, with an average Cα-root mean squared deviation of the top models (〈*C*α - *RMSD*〉) of 1.9Å and an average template modeling score of 0.71, which is very close to the optimum achievable with our simulation protocol (using contacts, secondary structure elements and beta sheet interactions from the reference structure, 〈*C*α - *RMSD*〉 = 1.4Å and TM score = 0.8); and the top evaluated *combined score* structural model has a *C*α - *RMSD* of only 1.5Å (Figure 4F). Consistent with somewhat lower precision of contact and secondary structure predictions, models generated with restrains from *epistasis* or *association scores* have on average a lower accuracy (〈*C*α - *RMSD*〉 = 3.4Å and 〈*C*α - *RMSD*〉 = 2.6Å, respectively), with *association score* models performing consistently better (Figures 4E and Extended Data 4F).

Together, this shows that deep mutation scanning alone is sufficient to accurately determine the structure of a protein domain.

### Contact prediction in additional protein domains

To test the generality of our approach, we analyzed two additional, incomplete deep mutational scanning datasets. First, a mutational scan of the 75 amino acid Pab1 RRM2 domain (Figure 5A), for which fitness was assessed in a complementation assay ^10^. Second, a mutational scan of the hYAP65 WW domain (Figure 5C), in which 33 out of 50 amino acids were mutated and fitness was assayed by binding to a polyproline peptide ligand in a phage display assay ^46^. Both datasets were created by ‘doped’ oligonucleotide synthesis and thus consist primarily of amino acid changes elicited by just one nucleotide change, which results in only 10% of possible double mutants being present. Additionally, their selection assays have smaller measurement ranges than that of the GB1 domain, which results in higher relative errors of fitness estimates as well as in negative epistasis being quantifiable for a smaller fraction of double mutants, as low as 0.8% in the case of the WW domain (Extended Data Figure 5A, see Table 1 for comparison of dataset properties).

**Figure 5.**
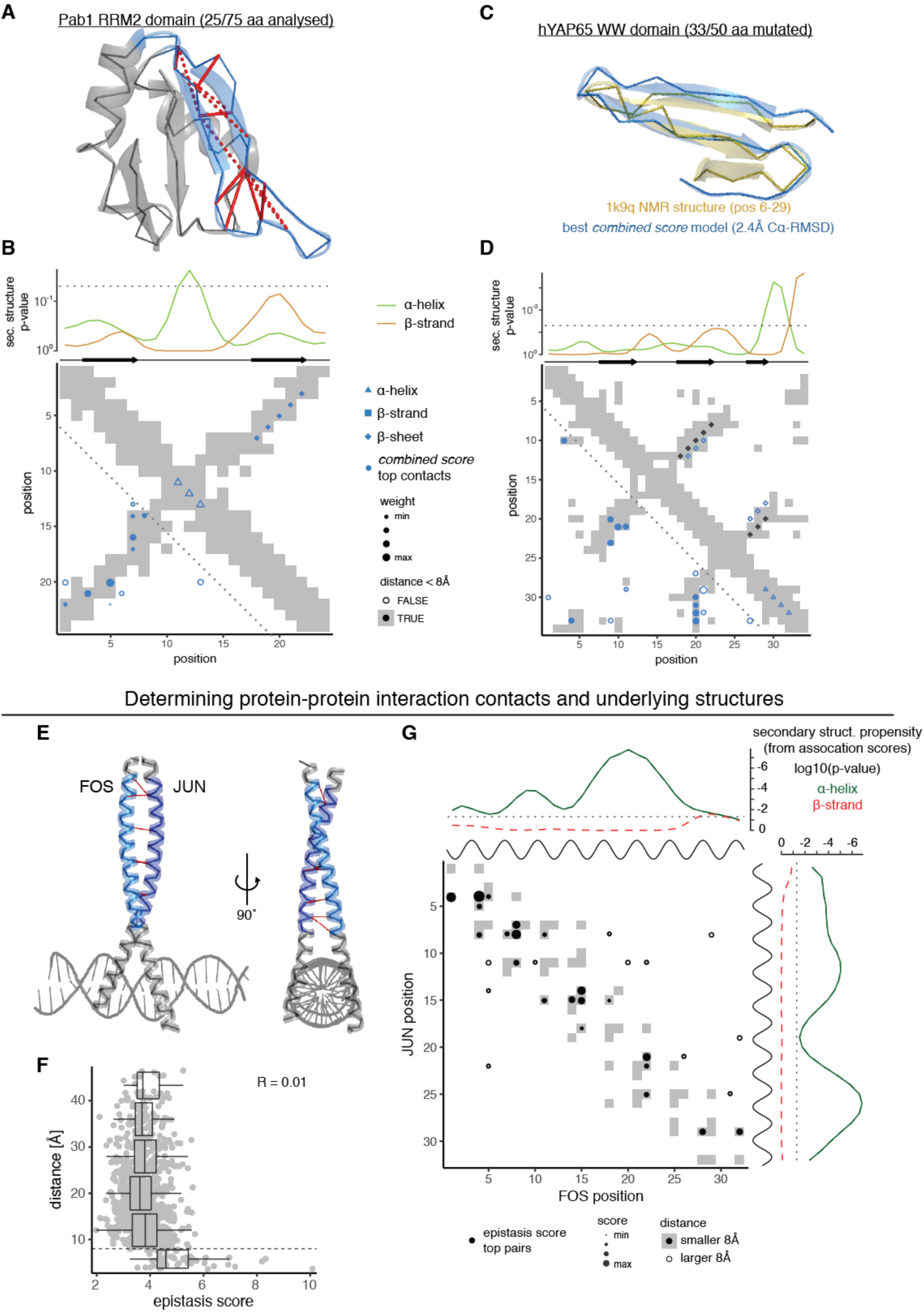
– Predicting structural contacts in two additional proteins and a protein-protein interaction. A. Pab1 RRM2 domain structure (PDB entry 1cvj) with 25/75 positions analyzed here highlighted in blue. Top 12 *combined score* position pairs are connected with red lines, solid if distance < 8Å, dashed otherwise. B. Structural predictions derived from *combined scores* in RRM domain. Upper plot shows secondary structure propensities from kernel smoothing (p = 0.05 indicated as dashed line). Just below are shown the secondary structure elements in the reference structure. Map shows top 12 *combined score* position pairs in lower left and secondary structure predictions in upper right triangle. Shape indicates type of prediction, fill indicates correct prediction. Underlying is the contact map of the reference structure (grey if < 8Å). C. Overlay of top structural model of hYAP65 WW domain (positions 6-29) generated with restraints from *combined score* (blue) and solution NMR structure (gold, PDB entry 1k9q). D. Structural predictions derived from *combined scores* in WW domain. Upper plot shows secondary structure propensities from kernel smoothing (p = 0.05 indicated as dashed line). Just below are shown the secondary structure elements in the reference structure. Map shows top 17 *combined score* position pairs in lower left and secondary structure predictions in upper right triangle. Shape indicates type of prediction, fill indicates correct prediction. Underlying is the contact map of the reference structure. Black diamonds indicate positions of beta sheet pairing in reference structure. Crystal structure of the leucine zipper domains of FOS and JUN with a DNA strand (PDB entry 1fos). The mutated regions (32 amino acids each) are highlighted in light blue (FOS) and dark blue (JUN). Top 10 epistasis score pairs are shown with red dashes. E. Distance of position pairs as a function of interaction scores. Boxplots are spaced in distance intervals of 8Å. Dashed horizontal line indicates 8Å. Pearson correlation coefficient is indicated. F. FOS-JUN trans interaction score map for top 32 position pairs with highest *epistasis scores*. Note that protein-protein interaction maps are not symmetric. Dot size indicates relative score; dot fill indicates distance below 8Å; underlying in grey is the contact map of the reference structure (PDB entry 1fos, distance < 8Å). Shown on top and to the right of the contact map are the known alpha helices and secondary structure propensities derived from *association scores* of FOS and JUN, respectively (black – known alpha helix; green – predicted alpha helix propensity, orange - predicted beta strand propensity; see Extended Data Figures 5F,G).

For the RRM domain, three 25 amino acid segments were mutated separately and we restricted our analysis to the central one, as it is the only segment that exhibits a reasonable number of intra-segment contacts in the reference structure (Figure 5A). We find that predicted tertiary contacts fall on or very close to known contacts in the region of the anti-parallel beta sheet and the intervening loop region (Figure 5B), with a precision of 57% for the top L/2 and 50% for the top L position pairs of the *combined score* (3-fold and 2.7-fold over expectation, respectively). Predicted beta strand propensities peak at the correct positions, albeit with low statistical significance; additionally, an alpha helical propensity is detected in the intervening loop region. Nonetheless, the correct anti-parallel beta sheet conformation at the correct position pairs is predicted.

For the WW domain, we find that top predicted tertiary contacts fall on or very close to known interactions between the beta strands and the N-terminal and C-terminal loop regions (Figure 5D), with a precision of 59% for the top L/2 and 38% for the top L position pairs of the *combined score* (3.9-fold and 2.5-fold over expectation, respectively). Secondary structure elements are not well predicted, but beta sheet interactions are predicted in the right anti-parallel conformation of *β*1 - *β*2 - *β*3, though the exact pairing between positions is off by one to two positions. We determined the three dimensional structure of the secondary structure-rich central part of the domain (positions 6 to 29, 24 amino acids), using restraints derived from top *combined score* pairs and PSIPRED-predicted secondary structure elements (see Methods). The top 5% of structural models have an average accuracy of 3.3Å 〈*C*α - *RMSD*〉 compared to the reference structure, which is on par with simulations using a set of ‘true’ contacts (*C*α - *RMSD* = 3.6Å) (Extended Data Figure 5C). Moreover, the structural model with the best XPLOR-NIH energy has an accuracy of 2.4Å *C*α - *RMSD* (or 2.0Å over 22 of the 24 residues) (Figure 5C). Despite similar precision of predicted contacts, *association* and *combined score*- derived WW domain structural models are more accurate than *epistasis score*-derived models (Extended Data Figure 5C).

Together these results strongly support the generality of our approach for extracting structural information from deep mutagenesis data, including from sparser and lower quality data.

### Contact prediction in a protein-protein interaction

Genetic interactions do not only occur between mutations within individual proteins but also between molecules that physically interact ^2^. We investigated a deep mutational scanning dataset of the coiled-coil interaction between the proteins encoded by the proto-oncogenes *FOS* and *JUN* (Figure 5E) ^9^. In this experiment, all possible single amino acid changes were made in each of 32 positions of each protein and the physical interaction of all single and (trans-)double mutants was quantified using a deep sequencing-based protein complementation assay. After filtering, the dataset contains 43% of all possible double mutants and has a median relative error of fitness measurements of 3.6% (Table 1).

When assessing the enrichment of epistatic interactions between positions in the two interaction partners we find a striking all-or-none relationship between *epistasis scores* and pair-distances (Figure 5F), with all distant pairs contained in a low *epistasis score* peak and only proximal interactions enriched for high *epistasis scores* (Pearson correlation coefficient R = 0.01, n = 1024). Indeed, the top 11 *epistasis score* pairs are all proximal interactions, and the precision of contact prediction is 75% for the top L/2 contacts and 66% for the top L contacts (12-fold and 10.5-fold over expectation). Moreover, top *epistasis score* pairs are evenly distributed across the interaction surface (Figures 5E and 5G).

When correlating epistatic patterns between columns of the epistatic enrichment matrices, one is comparing the epistatic interactions that two positions in FOS have with all positions in JUN. Therefore, the similarity of column-wise epistatic patterns reveals the *cis* relationships between positions in FOS (Extended Data Figure 5F). Similarly, correlating epistasis patterns across combinations of rows of the epistatic enrichment matrices reveals *cis* relationships between positions in JUN. We find that cis-interaction maps from *association scores* for both FOS and JUN are highly enriched for strong local interactions with an alpha helical periodicity; and applying our secondary structure prediction algorithms to the *cis*-interaction maps reveals strong alpha helix propensities across the full lengths of both FOS and JUN (Figures 5G and Extended Data Figure 5G).

This shows that deep mutagenesis of protein interaction partners can accurately predict direct contact across the interaction surface as well as the underlying structures of the interaction partners themselves.

### Deep learning improves contact prediction

Evolutionary coupling-based structural predictions have been successfully improved by machine learning approaches that transform the two-dimensional interaction score maps after learning the stereotypical patterns between evolutionary coupling-predicted contact maps and the actual contact maps of the known structures ^47,48^.

We tested whether such an approach can also improve deep mutagenesis-derived contact predictions. We applied a convolutional neural network approach called *DeepContact*, developed by Liu, et al. ^47^. The basic *DeepContact* architecture takes as a sole input a two- dimensional interaction score map that it then transforms based on the structural patterns it has previously learned on evolutionary coupling-derived contact predictions for representative families of the SCOPe database ^49^ (Figure 6A and Methods). When transforming evolutionary coupling-derived contact predictions of proteins not contained in the training set, this basic *DeepContact* architecture has been shown to improve contact prediction precision by about 10-20% ^47^.

**Figure 6:**
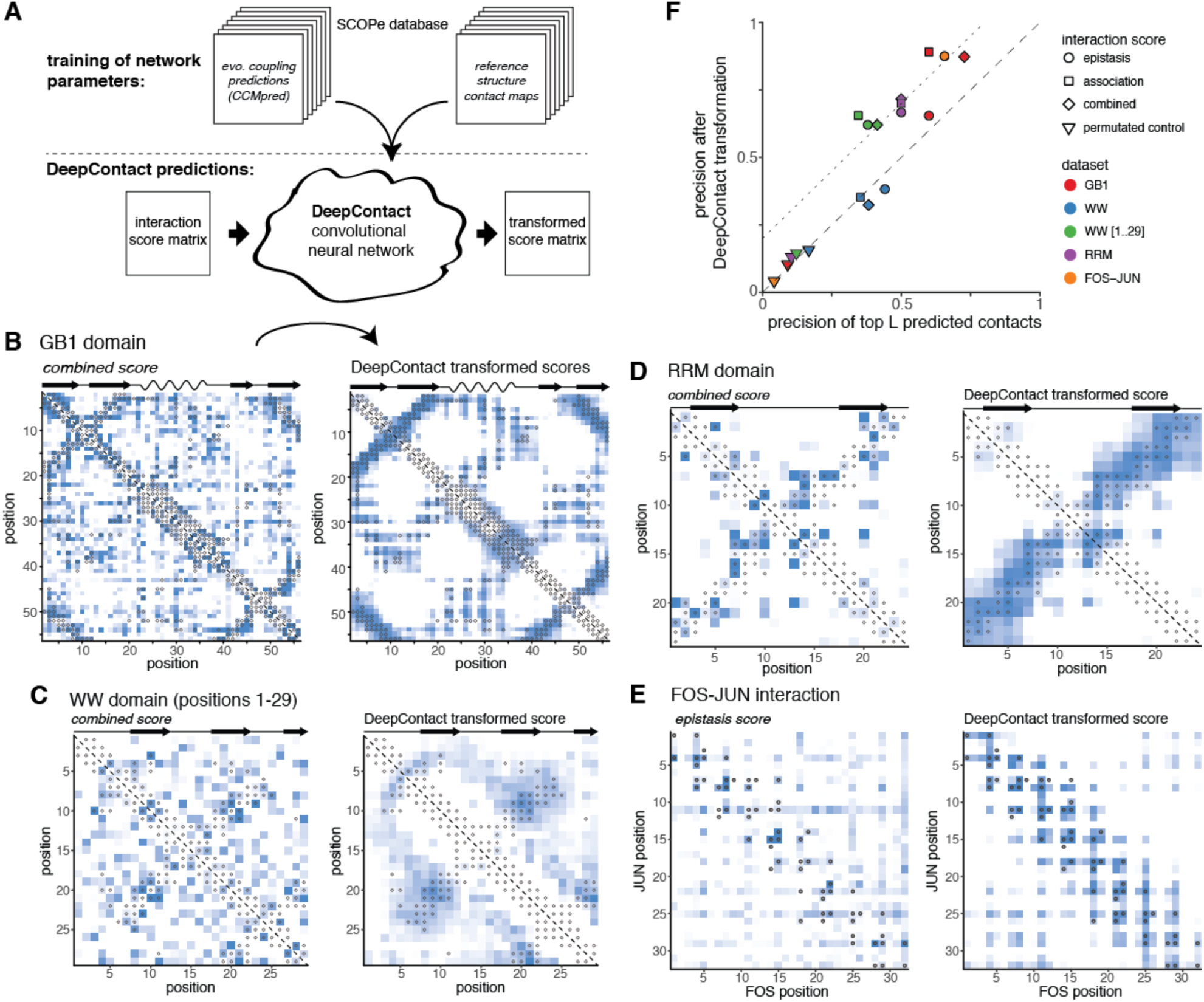
Deep learning improves contact prediction from deep mutagenesis data. A. DeepContact convolutional neural network transforms DMS-derived interaction score maps based on learned structural patterns ^47^. The particular DeepContact architecture used here takes as only input the DMS-derived interaction score map and transforms it based on structural patterns previously learned on an orthogonal and independent training set (in which it compared evolutionary coupling-derived contact predictions with contacts in known structures of representative protein families in the SCOPe database). B. GB1 domain *combined score* interaction map before (left panel) and after (right panel) transformation with *DeepContact* convolutional neural network. Heat maps show scores (normalized to have similar range). Grey open circles show contacts (side-chain heavy atom distance < 8Å) in reference structure. C. WW domain *combined score* interaction map before (left) and after (right) DeepContact transformation. Note that the maps shown here lack the 5 c-terminal positions (see Extended Data Figure 6B for full map). D. RRM domain *combined score* interaction map before (left) and after (right) *DeepContact* transformation. E. FOS-JUN trans-interaction *epistasis score* interaction map before (left) and after (right) *DeepContact* transformation. F. Precision of top L predicted contacts of different interaction scores before and after *DeepContact* transformation for the four datasets. Color indicates dataset, shape indicates interaction score. Permutated control score is average over three random permutations of *combined score* matrices (in case of FOS-JUN *epistasis score* matrices). Dashed diagonal line indicates no changes in precision, dotted diagonal line shows precision improvement of 20% after DeepContact learning.

We first transformed the GB1 domain *combined score* interaction map with the DeepContact network (Figure 6B). These transformations take as sole input our deep mutational scanning-derived predictions and include no evolutionary coupling or otherwise-derived structural predictors for GB1. The scores on the transformed map are much less noisy, with high scores exclusively focused in areas of known contacts, especially those of secondary structure element interactions, and areas devoid of contacts showing homogenously low scores. Moreover, the transformed scores are highly correlated with pair-distances in the reference structure (Pearson correlation coefficient R = -0.68, p < 10^-6^, n = 1225, Extended Data Figure 6A). The precision of top predicted contacts improves from 82% to 96% for L/2 and from 73% to 87% for L predicted contacts (Figure 6F). *Epistasis score*-derived predictions improve by about 5%, while a*ssociation score*-derived predictions improve by 29% at L predicted contacts. In contrast, randomized interaction score maps show no changes in prediction performance over random expectation after transformation with DeepContact.

Interaction score maps for the other datasets show similar improvements to GB1 both in terms of cleaner interaction score maps that resemble the reference contact maps as well as increases in contact prediction precision of up to 30% (Figure 6 C-F).

This shows that machine learning can substantially improve contact map prediction from deep mutational scanning data, allowing the use of sparser and lower quality data for accurate prediction.

### Minimal data quality requirements for successful protein structure prediction

We further investigated how robust our prediction strategy is to changes in data quality by artificially down-sampling the GB1 domain dataset, thus assessing the minimal requirements for deep mutational scanning datasets to be useful for structure prediction.

First, we considered the sequencing read coverage. The GB1 domain dataset consists of about 600 million sequencing reads ^11^. We find that artificially down-sampling the sequencing read coverage of the dataset to 25% or 10% hardly affects the precision of predicted tertiary contacts (Figures 7A). Only when using just 2.5% of sequencing reads (15 million) does the precision of top L contacts drop below 50%.

**Figure 7:**
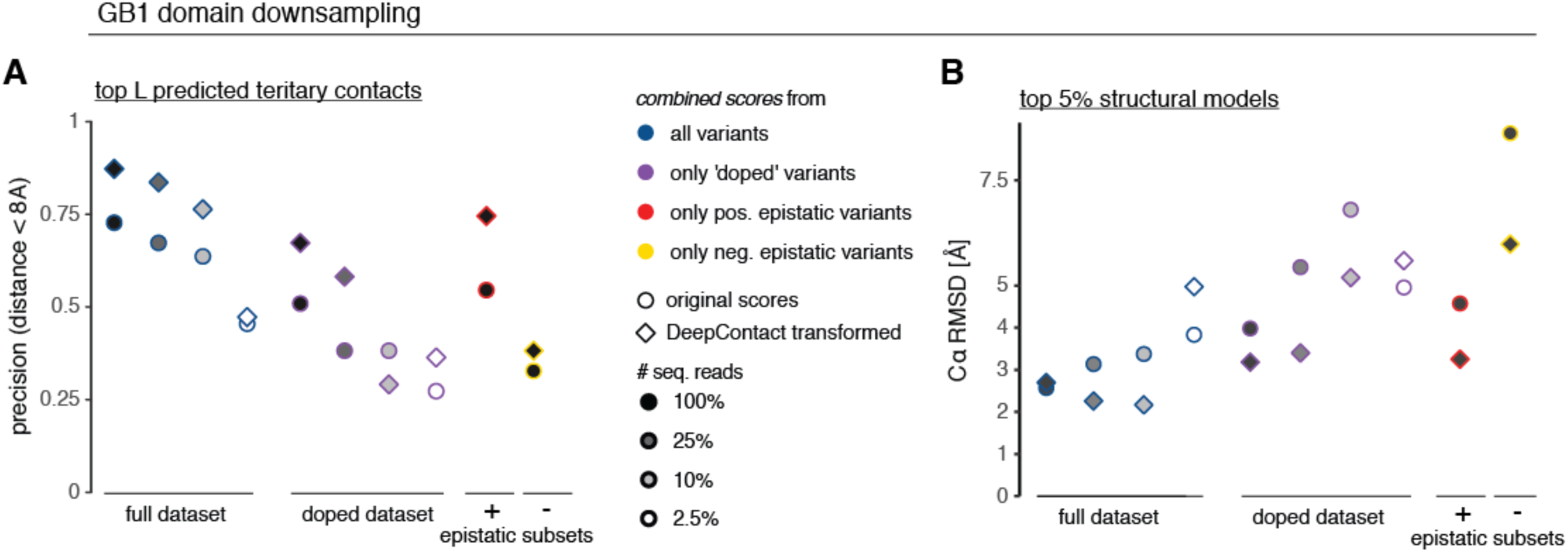
Deep learning allows contact and structure prediction from sparser and lower quality datasets. A. Precision of top L *combined score* position pairs for different down-sampled versions of GB1 dataset. Color indicates type of dataset (blue – full dataset, purple – ‘doped’ dataset, red – only positive epistasis information, yellow – only negative epistasis information), fill indicates number of sequencing reads used in analysis, shape indicates whether DeepContact has been used to transform the interaction score matrix. B. Accuracy 〈*C*α - *RMSD*〉 of top 5% structural models derived with tertiary contact restraints from down-sampled GB1 datasets compared to reference structure. Note that for better comparability, for these structural simulations only distance restraints were derived from *combined scores* but the same secondary structure restraints predicted from PSIPRED and no beta sheet pairing restraints were used for all simulations. Colors and fills as in panel A.

Next, we simulated a ‘doped’ dataset, by only considering amino acid mutations that can be reached by one mutation in the nucleotide sequence - thus reducing the coverage of double mutants to ∽10% (similar to RRM and WW domain datasets). The doped dataset with full sequencing read coverage exhibits a drop in precision of predicted tertiary contacts of about 20%. Moreover, the doped dataset shows an increased sensitivity to lower sequencing read coverage.

We also tested the effect of reducing the signal-to-noise ratio (i.e. the measurement range of selection assay relative to the median error of fitness estimates), which results in non-quantifiably of negative epistasis (Extended Data Figures 2D-F and 5A). We thus tested how our prediction strategy performs on the GB1 domain dataset when only positive epistasis information is available; and find that it results in a drop of precision of about 20%, comparable to that observed for a doped dataset. In contrast, only using negative epistasis information resulted in a drop to ∽35% precision, as low as a doped dataset with 10% sequencing coverage.

We evaluated secondary structure prediction performance of the various down-sampled GB1 domain datasets. Beta strand and alpha helix predictions are hardly affected by lowered data quality or partial epistasis information (Extended Data Figure 7A). In contrast, precision and recall of beta sheet positional pairing is strongly affected by dataset quality, although often the correct overall conformation of beta sheets is still recovered (Extended Data Figure 7B).

We next tested whether DeepContact could also improve prediction performance on these down-sampled datasets. Similar to interaction scores derived from full datasets, DeepContact transformation of *combined scores* derived from down-sampled GB1 datasets improves the precision of predicted contacts by about 10-25% even for quite low quality datasets, i.e. the complete datasets with at least 10% read coverage, doped datasets with at least 25% read coverage or the dataset with only positive epistasis information (Figure 7A).

Finally, we evaluated how differences in prediction performance of tertiary contacts affect structural modeling. We find that changes in accuracy of the top structural models roughly scale with changes in contact prediction performance (Figure 7B). Down-sampling of sequencing reads in the complete dataset from 100% to 2.5% leads to a drop in accuracy from 2.5Å to 4Å 〈*C*α - *RMSD*〉, which is roughly also the accuracy of top structural models from the doped dataset and the dataset using only positive epistasis information. Accuracies for lower quality datasets range from 5Å to 9Å 〈*C*α - *RMSD*〉. DeepContact increases the accuracy of the top structural models by up to 2.6Å. For the complete datasets with only 25% or 10% of sequencing reads, the top structural models have better accuracy than those from the complete dataset with full sequencing read coverage but untransformed scores. Also, structural models based on DeepContact transformed scores from the doped dataset with full or 25% sequencing coverage and those from the dataset using only positive epistasis information reach average accuracies of 3.2Å 〈*C*α - *RMSD*〉. Only for the two datasets with 2.5% sequencing read coverage do structural simulations based on DeepContact transformed scores not improve model accuracy.

Together these findings show that contact and structure prediction from deep mutational scanning data can also work for lower quality datasets and that the use of deep learning allows the use of much sparser and lower quality datasets.

## Discussion

We have shown here that simply quantifying the activity of a large number of single and double mutant variants of a macromolecule can provide enough information to determine its high-resolution 3D structure.

We found that although most epistasis within a protein occurs between positions that are not direct structural contacts, aggregation on position pairs, merging of positive and negative epistasis information and partial correlation analysis of epistasis patterns can successfully discriminate direct from indirect structural contacts. Thus, mostly indirect epistatic couplings can be transformed to predict accurate structural contacts and elements. We have shown that this approach works robustly across multiple protein domains and a protein interaction. Moreover, we have demonstrated that the application of a convolutional neural network previously trained on patterns of co-evolution in proteins of known structure both improves structure prediction and allows the use of much lower quality deep mutation datasets.

Determining structures by mutagenesis requires an *in vitro* or *in vivo* selection assay. For many important molecules and drug targets, specific selection assays based on known functions or interaction partners already exist ^11,14,15,46,50-55^. Additionally, many generic selection assays have recently been developed that should allow the stability or functional activity of many proteins to be assayed *in vivo* without the need for much prior knowledge about the protein under investigation ^35-38^. Moreover, many molecules have known interaction partners – proteins, DNA, RNA, or small ligands – for which *cis*- and *trans*-epistasis can thus be assessed by binding assays ^9,55^. We have shown here how *trans*-epistasis, for which library design is relatively easy, can lead to information about direct contacts in interaction surfaces as well as in the individual molecules.

Although structural information exists in the epistasis maps, our analyses and previous work ^5-16^ has shown that many epistatic interactions occur between positions that are not in direct structural contact. Indeed, in the GB1 domain the interactions are strikingly modular, with two mutually exclusive clusters of positive and negative epistatic interactions. This is consistent with many interactions being due to functional or energetic couplings between positions. The cluster of mostly positive epistatic interactions corresponds to a dynamic region involved in IgG binding ^11^. In contrast, the cluster of negative epistatic interactions identifies positions important for the thermodynamic stability of the domain ^56^, and the periodicity of local negative epistatic interactions provide evidence for a shift towards an alternative three-helical conformation that has been previously reported for this sequence family (Extended Data Figure 4E) ^57^. This modular organization of epistasis is thus reminiscent of the concept of energetically-coupled protein sectors identified from patterns of sequence co-evolution ^20,21,58^.

For macromolecules with very large numbers of homologs available in sequence databases, correlated changes in sequence can provide sufficient information for structure determination ^25-34^. However, for many proteins and RNAs insufficient numbers of homologs are available, and for fast evolving, recently-evolved or *de novo* designed molecules this is a fundamental limitation ^26,32,37,39^. Moreover, co-evolutionary analysis provides information on the average structure across a large set of homologs, whereas it is easy to envisage how deep mutagenesis could be used to directly determine alternative conformations of macromolecules when they are performing particular selectable functions. The success of evolutionary coupling analysis for predicting the structures of diverse folds and macromolecules does, however, strongly support the generality of the approach outlined here. The demonstration that a deep learning approach previously trained on evolutionary couplings dramatically improves the prediction of contacts from deep mutagenesis data further supports this.

As a proof-of-principle we have shown that information from deep mutational scanning experiments alone is sufficient for accurate structure prediction. In practice, however, integration with other structural information is likely to further boost performance. As a first step, we used a deep learning approach, DeepContact, that was trained on evolutionary couplings to learn stereotypical structural patterns in contact maps ^47^. DeepContact improved DMS-derived contact prediction precision by up to 30% for individual proteins (GB1, WW and RRM domains). Moreover, even though it had only been trained on data from individual proteins, it also improved DMS-derived contact predictions for the FOS-JUN protein-protein interaction. Integration with other structural predictors ^47,59,60^ and homology-driven structure modeling ^61,62^ is likely to further improve accuracy and lower the data quality requirements for structure determination by deep mutagenesis.

An analysis of incomplete and down-sampled variants of the GB1 dataset suggests that a high signal-to-noise ratio (measurement range relative to experimental error of fitness estimates), which allows both positive and negative epistasis to be quantified, is an important factor for generating datasets with a quality sufficient for protein structure prediction. For datasets with complete epistasis information, however, sequencing coverage hardly affected prediction performance. In contrast, prediction performance on the incomplete, ‘doped’ dataset was sensitive to sequencing coverage. Down to 25% sequencing coverage, however, performance could be recovered by deep learning. Together, these analyses suggest that our approach should be easily applicable to longer molecules. For example, with the experimental effort undertaken to create and sequence the 55 amino acid GB1 library, a protein of length ∽350 amino acids should be assayable at similar prediction performance (using a doped library with 25% sequencing coverage, i.e. 2.5% of the data: 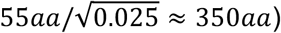. Such libraries for longer proteins could be created via fragment-based ligation approaches ^52^ or via random mutagenesis and barcode-variant linking ^35^.

Taken together, the results presented here establish deep mutagenesis as a new experimental strategy for structure determination. The approach that we have outlined is not the only one that can be envisaged to predict direct structural contacts from deep mutagenesis data, and other related approaches are also likely to work ^63^. The determination of macromolecular structures by physical techniques requires access to very expensive scientific infrastructure. In contrast, deep mutagenesis only requires techniques familiar to many molecular biologists and access to sequencing that is increasingly low cost and available to all. Most importantly, however, deep mutagenesis allows the structures of macromolecules to be studied whilst they are performing particular functions *in vitro* as well as *in vivo* in the cell. As such, deep mutagenesis opens up the possibility of low cost and high throughput determination of *in vivo* macromolecular structures by many molecular biology and genomics labs. A large-scale project to systematically determine the structures of proteins and protein domains should therefore be possible using the existing infrastructure of genomics institutes.

## Acknowledgements

We are grateful to Yang Liu and Jian Peng for making their DeepContact code available and for their advice. We thank members of the Lehner lab, T. Gross, G. Mönke, M. Bolognesi and C. Camilloni for discussions and feedback. This work was supported by a European Research Council (ERC) Consolidator grant (616434), the Spanish Ministry of Economy and Competitiveness (BFU2011-26206 and SEV-2012-0208), the AXA Research Fund, the Bettencourt Schueller Foundation, Agencia de Gestio d’Ajuts Universitaris i de Recerca (AGAUR, SGR-831), the EMBL-CRG Systems Biology Program, and the CERCA Program/Generalitat de Catalunya. J.M.S. was supported by an EMBO Long-Term Fellowship (ALTF 857-2016). This project has received funding from the European Union’s Horizon 2020 research and innovation programme under the Marie Sklodowska-Curie grant agreement No 752809 (J.M.S).

## Author Contributions

Conceptualization, B.L. and J.M.S.; Methodology, J.M.S.; Investigation, J.M.S.; Writing, J.M.S. and B.L.; Supervision, B.L.

## Additional Information

**Correspondence and requests for materials** should be addressed to B.L.

## Methods

### Datasets and preprocessing

#### Protein G B1 domain

Protein G B1 domain (GB1) deep mutational scanning data was obtained from the supplementary information of Olson, et al. ^11^. The data consists of summed read counts of three replicate experiments assaying the binding affinity of GB1 variants to immunoglobulin G (IgG).

Read frequencies of each single or double mutant variant in input library and output library (after binding affinity assay) were calculated as variant read counts relative to wild-type variant read counts. A variant’s fitness was calculated as the natural logarithm of the ratio of output to input or read frequency, i.e. 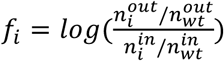, with n as read counts, superscripts denoting input or output sequencing library and subscripts denoting variant *i* or wild-type variant.

The standard error of fitness estimates was calculated from read counts under Poissonian assumptions, i.e. 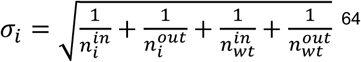 We note that this is a lower bound estimate of the actual error, due to the lack of replicate information.

Each measurement assay has a lower measurement limit due to unspecific background effects (Extended Data Figure 2A). In the case of the IgG-binding assay for GB1, this is presumably mainly due to unspecific carry-over on beads ^11^. The fitness values derived from the measurement are therefore a convolution of the actual binding affinities to IgG and nonspecific carryover, i.e.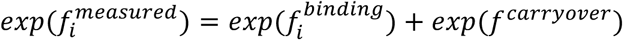 Fitness values of variants close to the lower measurement limit of the assay are therefore dominated by unspecific carryover effects. The lower measurement limit of the assay was estimated by two approaches that yielded identical estimates. One, from a kernel density estimate of the single mutant fitness distribution (R function *density* with parameter *bw* set to 0.15), where the position of the lower mode of the data corresponded to *f*^*carryover*^ = -5.85 (∽0.3% on linear scale). Two, from examining the fitness distribution of double mutants with expected fitness lower than -8 log-units, i.e. double mutants resulting from two lethal or nearly lethal single mutant variants, whose fitness values are thus expected to be dominated by background effects. The median of this background fitness distribution yielded an estimate of *f*^*carryover*^ = -5.85.

7% of double mutant variants were discarded due to too low sequencing coverage in input or output libraries (Extended Data Figure 2B). That is, variants that had less than 200 input reads and no output reads were discarded, because it is not possible to determine their fitness. Above 200 input reads, variants without output reads are certain to be dominated by nonspecific carryover effects. These variants were retained and their fitness was calculated by setting their output read count to 0.5.

### GB1 down-sampling

Down-sampling of the full GB1 dataset was performed in three different ways.

For the ‘doped’ datasets, we only allowed amino acid changes created by one nucleotide mutation from the wild-type sequence (ENA entry M12825). To down-sample the sequencing read coverage, for each variant we picked as a down-sampled read count the draw of successes from a binomial distribution with the number of sequencing reads in the full datasets as trials and the target down-sampling rate (25%, 10% or 2.5%) as chance of success. For the read down-sampled and doped datasets (and combinations of both), the analysis workflow for the full dataset was repeated.

For the down-sampled datasets taking only positive or negative epistatic information into account, we calculated *epistasis* and *association scores* from epistatic enrichment matrices and partial correlation matrices of only positive or negative epistasis information. Instead of merging positive and negative matrices and then calculating z-scores, we only calculated z-scores with the individual errors from positive or negative epistasis information. The *combined scores* (for which results are reported) for each set was then calculated as for the full dataset by summing standardized *epistasis* and *association scores*.

### hYAP WW domain

hYAP WW domain data was obtained from Sequence Read Archive (SRA) entry SRP015751 ^46^. Paired-end reads were merged with USearch ^65^ and merged reads with any base having a Phred base quality score below 20 were discarded. Read counts from the two technical sequencing replicates were merged and read counts for the same amino acid variants with at most one synonymous mutation in another codon were summed up. The dataset consists of an input library and three output libraries after consecutive rounds of selection in a phage display assay. Fitness was estimated as the slope of log frequency (variant counts divided by wild-type counts) changes over the rounds of selection experiment ^46^. For each variant at each selection step a Poissonian error of 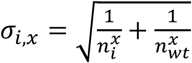was calculated, with x denoting the selection step.

Slopes were calculated as weighted straight line least square fits ^66^. Comparison of library-wide changes in variant frequencies between selection rounds suggested differential selection pressures across the rounds. We thus applied a non-equidistant spacing of 0.6, 1.17 and 1.22 between selection rounds when calculating slopes. Only variants that have more than 10 reads in the input library and at least one read after the first selection were retained for further analysis (45% of constructed double mutants). The lower fitness limit was calculated as the weighted mean fitness of all variants containing STOP codons (-0.78 in log-fitness units).

### Pab1 RRM2 domain

Pab1 RRM2 domain data was obtained from the Supplementary Table 5 of Melamed, et al. ^10^. Reported enrichment scores were log-transformed to obtain fitness values. Output reads per variant were deduced from the number of input reads times the enrichment score and used to calculate a Poissonian error of the fitness estimate. Single mutant count data is not provided and we thus estimated the error of single mutant fitness estimates to be 0.01. Lower bound of fitness assay was estimated as weighted mean fitness of all double mutant variants containing STOP codons (-3.1 log-fitness units).

### FOS-JUN interaction

Raw count tables were provided by Guillaume Diss ^9^. The dataset consists of input and output sequencing libraries after selection for physical interaction between the two proteins in a protein complementation assay in three biological replicates. Per sequencing library, read counts from all synonymous variants were summed up. Only variants that had more than 10 reads in each of the three input libraries were used for further analysis (43% of double mutants). Per input/output replicate, fitness of each variant was calculated as the log change in frequency compared to the wild-type variant (as for GB1). A Poissonian error for each variant’s fitness estimate was derived.

The dataset has a large dynamic range, thus many low-fitness variants with low input read coverage have very low or no output read counts (per replicate ∽1/3 of variants have below 3 output counts, ∽15% of variants have zero output counts), effectively reducing the dynamic range of the assay for low input variants and distorting the estimate of the overall fitness distribution (see Extended Data Figure 5E). To overcome this, we implemented a Bayesian estimator of fitness. For each double mutant variant, we first identified the 1000 nearest neighbors in single mutant fitness space (i.e. those double mutants whose respective single mutant fitness values are similar to the single mutant fitness values of the variant under consideration) with sufficient input coverage (more than 100 reads in the input library). From this set of 1000 nearest neighbors we calculated the expected distribution of double mutant fitness values, which served as a prior distribution. For the variant under consideration we calculated the likelihood distribution of fitness values given its input and output read counts under Poissonian assumptions. Fitness was then estimated as the mean of the distribution resulting from the multiplication of prior and likelihood distributions. Error of fitness estimate was estimated as the standard deviation of the resulting distribution. Estimated fitness from the three replicate experiments were merged by weighted averaging.

### Epistasis classification

Epistasis was calculated from a non-parametric null model (Figure 1B) in order to account for nonlinearities close to the lower limit of the fitness assay measurement range, non-specific epistatic behavior resulting from e.g. thermodynamic stability thresholds as well as differential uncertainty of fitness measurements across the fitness landscape, due to lower read counts in the output for low fitness variants.

First, double mutant fitness values were corrected by subtracting the average local fitness computed using a two-dimensional local polynomial regression (using R function *loess* with span = 0.2). This was necessary to avoid boundary effects of quantile-based fits in boundary regions with non-zero slopes. 5th and 95th percentile surfaces were then fit to these corrected double mutant fitness values, by computing for each double mutant variant the 5th and 95th percentile of the fitness distribution made up of the 1% closest neighbors in single mutant fitness space. Double mutant variants with fitness values below the 5th or above the 95th percentile were categorized as negative or positive epistatic, respectively (Figure 1B).

The evaluation of positive or negative epistasis was, however, restricted to specific subsets of the data where measurement errors do not impede epistasis classification (Extended Data Figure 2C). The data subset deemed suitable for positive epistasis classification is limited to regions where

- the 95th percentile fitness surface is below wild-type fitness
- at least one single mutant fitness value is significantly smaller than wild-type fitness at two standard errors
- the expected fitness (sum of both single mutant fitness values) is not significantly higher than wild-type at two standard errors

The rationale for these criteria is to avoid double mutants from two neutral single mutants, because these are dominated by measurement noise of overabundant wild-type like variants. No restrictions were instead applied to the lower limits of the measurement range, because otherwise no/little epistasis quantification would have been available for several positions with very strong detrimental effects as well as because strong positive epistatic effects are observed in these regions, despite the dominance of background measurement effects.

The data subset in which variants were potentially classified as negative epistatic is limited to data regions where

- the 5th percentile fitness surface is above the 95th percentile of the background effect distribution; this value is derived from the 95th percentile of double mutant fitness values with expected fitness below -8 (analogous to lower fitness limit estimation, see above).
- both single mutant fitness values are significantly higher than the lower limit of the fitness assay measurement range at two standard errors
- the expected fitness (sum of both single mutant fitness values) is not significantly higher than wild-type at two standard errorss

The rationale for criteria 1 and 2 is to avoid background measurement effects that make negative epistasis quantification unreliable.

As a result of these restrictions as well as differences in initial coverage, the number of double mutant variants that can be used to assess positive and negative epistasis varies substantially across position pairs and datasets (see Table 1, Extended Data Figures 2D-F and 5A&D).

### Epistatic interactions (*epistasis scores*)

We derived several interaction scores to estimate which position pairs are in close contact in the tertiary structure (see Figures 2A and 3A,B and Extended Data Figure 1). These scores are based on summarizing epistasis information on the position pair-level and accounting for the uncertainty inherent in the summarized estimates due to differential error of fitness estimates across the measurement range as well as varying numbers of double mutants amenable to epistasis classification (see Table 1, Extended Data Figures 2D-F and 5A&D).

To summarize epistasis information on the position pair-level, we calculated the fraction of positive or negative epistatic variants per position pair. The fraction of positive epistatic variants per position pair is the number of positive epistatic variants divided by the number of all variants that lie in the double mutant space amenable to positive epistasis classification (Extended Data Figure 1, step 5b, equivalent calculation for negative epistasis fraction). Because enrichments with positive and negative epistatic variants per position are anti-correlated (Extended Data Figure 3A), we treated both separately and only aggregated them to derive the final interaction scores.

To estimate the uncertainty in epistatic fractions per position pair for downstream analyses we implemented a re-sampling approach (Extended Data Figure 1, step 5, described here for positive epistatic variants, but equivalent for negative epistatic variants). In each of 10.000 re-sampling runs:

- each variant’s fitness was drawn from a normal distribution with the measured fitness as mean and the uncertainty due to sequencing coverage as standard deviation 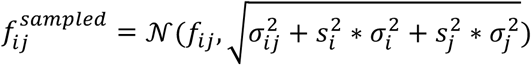, with *s*_*x*_ as the local slope of the median fitness landscape in direction of the respective single mutant (step 5a)
- positive epistasis of variants was res classified given the drawn fitness values (also step 5a)
- each position pair’s fraction of positive epistatic variants was drawn from the posterior probability distribution of how likely an underlying true fraction of epistatic variants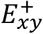.Is to generate the observed fraction of epistatic variants given the finite number of overall variants, i.e.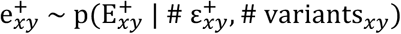 (step 5b). The posterior probability distribution is the product of a prior probability distribution – the kernel density estimate of the expected epistatic fractions across all position pairs (calculated using R function *density* with parameter *bw* set to 0.05) – and the likelihood function for the underlying true fraction of epistatic variants given the observed fraction of epistatic variants and the overall number of variants under binomial sampling assumptions

To derive an interaction score from the epistatic fractions per position pair, mean positive and negative epistatic fractions across resampling runs were combined by weighted averaging, with weights as the inverse variances of epistatic fractions across resampling runs, i.e.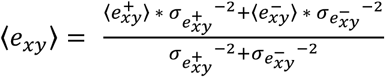

To arrive at the final *epistasis score*, the mean epistatic fractions were further normalized by their combined uncertainty,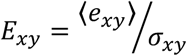,with 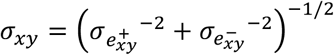(step 6).

### Epistasis pattern correlations (***association scores*)**

In addition to the *epistasis score* we derived an interaction score from the partial correlation of epistasis patterns between position pairs, termed *association score*. The rationale behind this score is that proximal positions in the protein should have similar distances and geometrical arrangements towards all other positions in the protein and should therefore also have similar patterns of epistatic interactions with all other positions.

In each re-sampling run we constructed a symmetric matrix of the drawn positive epistatic fractions 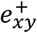.with number of rows and columns as the number of mutated positions (Extended Data Figure 1, step 5c, equivalent for negative epistatic fractions). Missing values (positions pairs without observed variants) were imputed by drawing a random value from the overall distribution of epistatic fractions. A pseudo count equal to the first quartile of the epistatic fraction distribution was added to each epistatic fraction. Diagonal elements (epistatic fractions of a position with itself) were set to 1. The matrix values were transformed by the natural logarithm (to make distributions more symmetric, thus correlations are not dominated by few position pairs with large epistatic fractions) and for each pair of columns the Pearson correlation coefficient was calculated to arrive at the correlation matrix (step 5d).

A shrinkage approach was used to improve the estimate of the correlation matrix ^67^. In brief, the empirical correlation matrix is shrunk towards the identity matrix in order to minimize the mean-squared error between estimated and true correlation matrix. Additionally, this yields a positive definite and well-conditioned correlation matrix, suitable for inversion. All computations on correlation matrices, shrinkage and matrix inversion were performed with the R package *corpcor*, functions *cor.shrink* and *pcor.shrink* ^67^.

Partial correlations of epistatic patterns between each position pair were calculated by inverting the shrunk correlation matrix and normalizing each off-diagonal entry of the inverted matrix by the geometric mean of the two respective diagonal entries, i.e.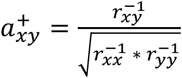, with.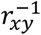.as the (x,y)-entry of the inverted correlation matrix (Extended Data Figure 1, step 5d). Equivalent to the *epistasis score*, positive and negative partial correlation estimates were merged by calculating weighted averages of their mean estimates across re-sampling runs, with weights as the inverse variances across resampling runs, i.e.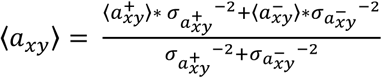,and the final *association score* normalized by the combined uncertainty,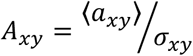 with 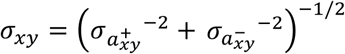.(step 6).

### Aggregating *epistasis* and *association scores* (*combined scores*)

We further derived a *combined score* by summing the standardized *epistasis* and *association scores*, i.e 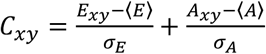. We note that this is a naïve approach to combining the information from these two complementary sources, and surely more sophisticated approaches that further improve proximity estimates can be developed.

### Secondary structure prediction

We used a two-dimensional kernel smoothing approach to predict secondary structure elements from interaction score matrices (Figure 4A-C). For a given position along the linear chain (on the diagonal of the interaction score matrix), interaction scores (off the diagonal) are integrated with distance-specific weighting according to the kernel, which reflects the known geometry of secondary structures.

The alpha kernel takes on a sinusoidal profile perpendicular to the diagonal to weight interactions according to whether the position pair considered should have congruent side-chain orientations (see diagonal and perpendicular profiles in Figure 4B). The kernel was defined as 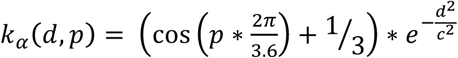,with *d=│2x-i-j│* as the diagonal distance of the interaction *ij* (off the diagonal) to the reference position *x* (on the diagonal) and *p = │i - j│* as the perpendicular distance of the interaction off the diagonal. The kernel weight for positions with p > 5 was set to 0, thus only including interactions across little more than the first helical turn. Finally, *c* = 4 is the integration scale for the Gaussian kernel along the diagonal. While smaller integration scales do yield nosier results and longer integration scales can lead to non-detection of shorter secondary structure stretches, we found that in practice our whole approach (including the actual detection algorithm described below) is robust to alterations of the integration length.

The kernel smoothed alpha value for a given position x on the diagonal is then calculated as the sum over all interaction scores times their kernel weights *K*_α,*x*_ =∑_*i*_ ∑_*j*_ *k*_*α*_ (*d, p*) *\* S*_*i j*_, where *S*_*i j*_ is one of the interaction scores (*epistasis, association* or *combined score*) at position pair *i j*.

The beta kernel takes an alternating profile perpendicular to the diagonal to weight interactions according to alternating side-chain orientations in a beta strand and was defined as 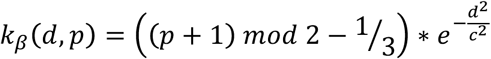, with *c* = 4. Only interactions with perpendicular distances equal or smaller than two (i.e. *k*_β_ (*d, p* > 3) = 0) were included.

To calculate whether kernel-weighted interaction scores of a specific position are larger than expected, they were compared to kernel-weighted scores obtained from 10^4^ randomized control datasets. Randomization was performed by shuffling all interaction scores, while preserving matrix symmetry, and kernel-weighted interaction scores from randomized control datasets were calculated for each position independently to control for possible boundary effects in positions close to the borders of the protein chain. A p-value for each position was calculated as the fraction of random controls with kernel smoothed values above that of the real data.

Secondary structure elements were identified by searching for continuous stretches of positions with high propensities to belong to either alpha helices or beta strands. The following workflow was implemented:

1. calculate a combined p-value for seeds of length 3 by combining position-wise p-values using Fisher’s method for both alpha and beta kernel smoothed interaction scores
2. separate positions according to whether combined p-values of seeds from alpha or beta kernels are more significant, i.e.
  2. 1. for alpha helical propensity calculations only consider stretches of at least 5 consecutive positions for which the combined p-value of seeds for alpha kernel smoothing is smaller than that from beta kernel smoothing (thus setting the lower size limit of alpha helical elements to five)
  2. 2. for beta strand propensity calculations only consider stretches of at least 3 consecutive positions for which the combined p-value of seeds for beta kernel smoothing is smaller than that from alpha kernel smoothing (thus setting the lower size limit of beta strands to three)
  For alpha helices and beta strands separately and while combined p-values of seeds < 0.05
3. select the most significant seed
4. test whether extension to any side yields lower combined p-value
  4. 1. if yes: extend seed in this direction and repeat step 4
  4. 2. else: repeat step 4 once to see whether further extension in same direction yields lower combined p-value
    4. 2. 1. if yes: extend and repeat step 4
    4. 2. 2. else: proceed to step 5
5. fix as secondary structure element and delete all ‘used’ p-values (and combined seed p-values), such that other elements cannot incorporate them
6. check whether other already fixed elements are adjacent or at most one position away if yes: merge both elements
7. repeat steps 3-6 until no more seeds with combined p-value < 0.05 are left

This yields a list of predicted locations of secondary structure elements. We note that the secondary structure elements predicted from deep mutational scanning data could be compared to and combined with predictions derived from other tools, such as PSIPRED (Jones, 1999), to further improve reliability.

To detect beta sheet interactions a modified beta strand kernel was used. In contrast to beta strand detection, the beta sheet interaction kernel is centered on each off-diagonal position. For beta sheet kernels diagonal and perpendicular distances are therefore modified as *d* = │*x* + *y* - *i* - *j*│ and *p* = │*x* - *i* - (*y* - *j*)│. The kernels to detect parallel and anti-parallel beta sheets differ in which is their ‘diagonal’ direction, i.e. the direction at which consecutive position pairs interact in the beta sheet (Extended Data Figure 4A). Therefore, parameters *d* and *p* were swapped for the anti-parallel beta sheet kernel. Also, because these positions can be deemed the most crucial for deciding whether a position participates in a beta sheet interaction or not, we up-weighted these positions (those with *p* = 0) in the kernel by a factor of two, i.e.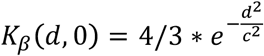.

Beta sheet interactions were identified by searching for the most significant stretches of parallel and anti-parallel interactions (similar to workflow for alpha helices and beta strands), then identifying the set of most significant interactions that is consistent with previously predicted secondary structure elements.

In particular, step 1 & 3-7 from the above-described workflow were performed for the parallel beta sheet kernel on each sub-diagonal (parallel to the main diagonal) of the interaction score matrix separately; and for the anti-parallel beta sheet kernel on each perpendicular diagonal of the interaction score matrix separately.

The steps were modified as follows:

- for anti-parallel beta sheet interactions, only positions with a distance greater than 1 from the main diagonal were used to calculate seed p-values (assuming anti-parallel beta sheet interactions need a turn of at least length two to be connected)
- for parallel beta sheet interactions, only sub-diagonals with a distance greater than 4 from the main diagonal were considered (assuming parallel beta sheet interactions of two adjacent beta strands need a linker region)

We extended the workflow with the following steps to predict beta sheet interactions within the protein domain:

8. compute association of seeds with known beta strands (e.g. seed positions overlap strand 1 on one side and coincide with strand 3 on the other side)
9. while there are seeds with p < 0.05: pick most significant seed from either the parallel or anti-parallel sheet subset
10. check consistency with secondary structure elements
  10. 1. discard the seed and jump back to step 9 if
    10. 1. 1. it is overlapping or too close to an alpha helix or the linker region between two beta strands that interact (minimal distance smaller one)
    10. 1. 2. at least one of the two strands it is associated with already has two other beta sheet interactions or the total number of beta sheet interactions exceeds 2*(#beta strands – 1)
  10. 2. modify secondary structure elements and start anew from step 3 if
    10. 2. 1. one side of the seed is not associated to a known beta strand: create this beta strand
    10. 2. 2. if both sides of the seed are associated with the same known strand: split the strand and create a linker region in-between the strands
  10. 3. else fix the beta sheet interaction and delete all other interactions that are associated with the same strands and haven’t been fixed yet, jump back to step 9
11. if no more seeds with p<0.001, finish
12. update beta strands: keep only those positions that are part of a beta sheet interaction

For beta sheet pairing detection in the GB1 domain (as reported in Figure 4D and Extended Data Figures 4A-C and 7B) we used as input the secondary structure element predictions derived from the deep mutational scanning data (as shown in Figure 4A-C and Extended Data Figure 4D). For the RRM and WW domains, we used as input PSIPRED predicted secondary structure elements, due to the insufficient signal from secondary structure element predictions from deep mutational scanning data.

### Protein distance metrics

The minimal side chain heavy atom distance, i.e. the minimal distance between any two side-chain heavy atoms of a position pair (in case of glycine, Cα), was used as the general distance measure. A direct contact was defined as minimal side-chain heavy atom distance < 8Å. For all evaluations of predicted contact precision we only considered position pairs with non-trivial tertiary contacts (those with a linear sequence separation of greater than 5 positions).

We do find that, while using all heavy atoms to calculate distances increase the true positive rates of predicted contacts by about 10%, side-chain heavy atom distances display much higher true positive rates over random expectation, thus suggesting that side-chain interactions are more informative for epistatic interactions (Extended Data Figure 7C).

The Floyd-Warshall algorithm (implemented as custom script in R) was used to calculate the minimal number of edges <8Å that connect any two positions in the protein.

Reference structures used as comparison were

- GB1 domain: PDB entry 1pga, X-ray diffraction structure ^68^
- WW domain: PDB entry 1k9q, solution NMR structure ^69^
- RRM domain: PDB entry 1cvj (chain A), X-ray diffraction structure of human Pab1 ^70^; note that the central section of the yeast RRM domain analysed is one nucleotide longer than the corresponding homologous region in the human RRM domain. We thus arbitrarily removed position 14 (in the loop region) when comparing the DMS-derived predictions to the human Pab1 structure.
- FOS-JUN interaction: PDB entry 1fos (chains E and F), X-ray diffraction structure ^71^ We found that precision or accuracy calculated against other reference structures differed only marginally, thus we have limited reporting to the aforementioned PDB entries.

### Protein folding

To *ab initio* determine protein structures, we performed simulated annealing using the XPLOR-NIH modeling suite ^45^ with structural restraints derived from the deep mutational scanning data. Simulations were performed in three stages, in each of which 500 structural models were generated. Stages 1 and 2 served to identify inconsistencies among defined structural restraints. Additionally, in stage 2 an average structure of the best 10% of models was calculated. Stage 3 served to refine this average structure to obtain a final set of best models.

Restraints from top predicted contacts (position pairs with highest interaction scores and linear chain separation greater than 5 positions) were implemented by setting Cβ-Cβ atom distances (Cα in case of Glycine) between positions to range between 0 and 8Å and weighting the restraints according to their relative interaction score (interaction score divided by mean interaction score of all predicted contacts used).

Restrains from secondary structures elements were implemented as dihedral angle restraints. Dihedral angles of both beta strands and alpha helices were set to range between values commonly observed in crystal structures ^72^, for alpha helices Φ_α_ = -63.5° ± 4.5° and ψ_α_ = -41.5° ± 5° and for beta strands Φ_β_ = -118° ± 10.7° and ψ_β_ = 134° ± 8.6°.

Restraints for beta sheet interactions were implemented by setting H-N:O=C hydrogen bond distances between interacting positions to range between 1.8 and 2.1Å ^72^, with weight one. Predictions of beta sheet interactions derived from deep mutational scanning data yield a string of interacting positions, but hydrogen bonding in beta sheets occurs in specific non-continuous patterns between position pairs (between alternating positions off the interaction diagonal in parallel beta sheets and between every second set of position pairs in anti-parallel beta sheets). Specifically, for each set of interacting positions there are two alternative patterns of hydrogen bonding possible. These alternative possibilities of pairing were implemented as mutually exclusive selection pairs with the “assign … or” syntax in Xplor-NIH.

Distance restraints were implemented in XPLOR-NIH as NOE (nuclear Overhausser effect) potential, with potential type set to “soft” for stages 1 and 2 and “hard” for the final simulation stage. Dihedral angle restraints were implemented via the CDIH potential.

After simulation stages 1 and 2 restraints were checked for their consistency with predicted structural models. First, structural models were clustered according to their violations of distance and dihedral angle restraints (k-means clustering, k = 4). Clusters were ranked by the mean total energy (from all energy potentials used) of their 50 models with lowest total energy (or all of their structures if clusters are smaller 50 models). From the 50 models with the lowest total energy from the top-ranked cluster (or however many top-ranked clusters were necessary to arrive at 50 models) the fraction of models that violate specific restraints was recorded. For the subsequent simulation stage, distance restraints were down-weighted according to the fraction of models that violated them, *w*_*x,i*_ = *w*_*x,i-1*_ *\** (1 – *f*_*x*_)^2^, and distance restraints with a weight below 0.1 were discarded. There is no option to weight dihedral angle restraints, thus instead dihedral angle restraints with a ‘weight’ below 1/3 were discarded for the subsequent simulation stages.

The top 5% structural models from simulation stage 3 were evaluated against the reference structure. The TM-score program (update 2016/03/23) was used to calculate the Cα root mean squared deviation and the template modeling score ^73^.

Several types of control simulations were performed to judge the predictive power of restraints derived from deep mutational scanning data. As a negative control we performed simulations without restraints from predicted contacts and beta sheet interactions, but with restraints from secondary structure elements predicted by PSIPRED (version 3.3, ^43^). As a positive control we performed simulations with restraints derived from the reference structure. Here, L true contacts of position pairs with linear chain distance greater than 5 amino acids were randomly sampled and beta sheet interactions were determined by PyMOL ^44^. These simulations serve as a positive control and give the maximally achievable accuracy of our Xplor-NIH workflow.

For the WW domain, simulations on the full mutated 33aa section gave mediocre results, both when using combined scores with PSIPRED predicted secondary structure (5.8Å Calpha-RMSD), as well as when using perfect information from the reference structure (4.1Å Calpha-RMSD). Upon inspection, this seemed to be an issue of the unstructured tail regions. We thus conducted structural simulations for a truncated version of the WW domain using only mutated positions 6-29 (the core region including the three beta strands).

For structural simulations of down-sampled GB1 datasets (and DeepContact transformed versions thereof) we used distance restraints derived from top predicted contacts and secondary structure restraints derived from PSIPRED predictions, but no restraints for beta sheet pairing. This was done to avoid skewed results due to false beta sheet pairing predictions in low quality datasets (Extended Data Figure 7B). For structural simulations from DeepContact-transformed predictions, we find that using more tertiary contacts results in better models. We conclude that this is because the deep learning algorithm focuses many strong predictions in few structural features (such as interactions of secondary structure elements), which are therefore the top contacts. Restraints in other regions of the protein are therefore only included if more predicted contacts are used for restraint calculations, therefore improving structural predictions. Because of this, when comparing structural simulations from scores derived before and after deep learning, we compare the top 5% of structural models derived with the top L predicted contacts from original scores with those derived with the top 1.5*L predicted contacts from DeepContact transformed scores.

### DeepContact learning

DeepContact software was obtained from GitHub (https://github.com/largelymfs/deepcontact) ^47^. We are grateful to Yang Liu and Jian Peng for also making - without any hesitation - their basic DeepContact network architecture available on their GitHub repository and helping us with the implementation. The DeepContact architecture used here only takes one 2D input of predicted contact scores and returns a 2D map of transformed scores (denoted as “DeepContact CCMPred only” in ^47^ and described in the first paragraph of the result section therein). The DeepContact architecture employed came with a pre-trained network model that had been trained on solved structures of the 40% homology filtered ASTRAL SCOPe 2.06 database (see GitHub repository and Liu, et al. ^47^), which were filtered to avoid structure and sequence redundancy of the training data. Because CCMpred scores ^74^ are distributed in the range of 0 to 1, we pre-normalized our deep mutational scanning derived interaction scores to this range (such that the minimum score on the interaction score matrix was 0 and the maximum score was 1) before providing them as an input to DeepContact. As negative control, we created for each dataset three random permutations of *combined score* matrices (while preserving matrix symmetry; in case of FOS-JUN dataset non-symmetric *epistasis score* matrices were permutated), which were transformed by the DeepContact algorithm. These control datasets show no increased precision of random expectation (Figure 6F).

### Code availability

Data was analyzed with custom scripts written and executed in R programming language, version 3.4.3. Structural simulations were performed with Xplor-NIH modeling suite version 2.46. Analysis scripts are available at https://github.com/lehner-lab/DMS2structure.

### Data availability

No primary data was generated in this study. Processed interaction scores for all datasets are included in Supplementary Table 1. All intermediate steps of data processing can be recapitulated with the scripts at https://github.com/lehner-lab/DMS2structure.

### Extended Data Figures

**Extended Data Figure 1:**
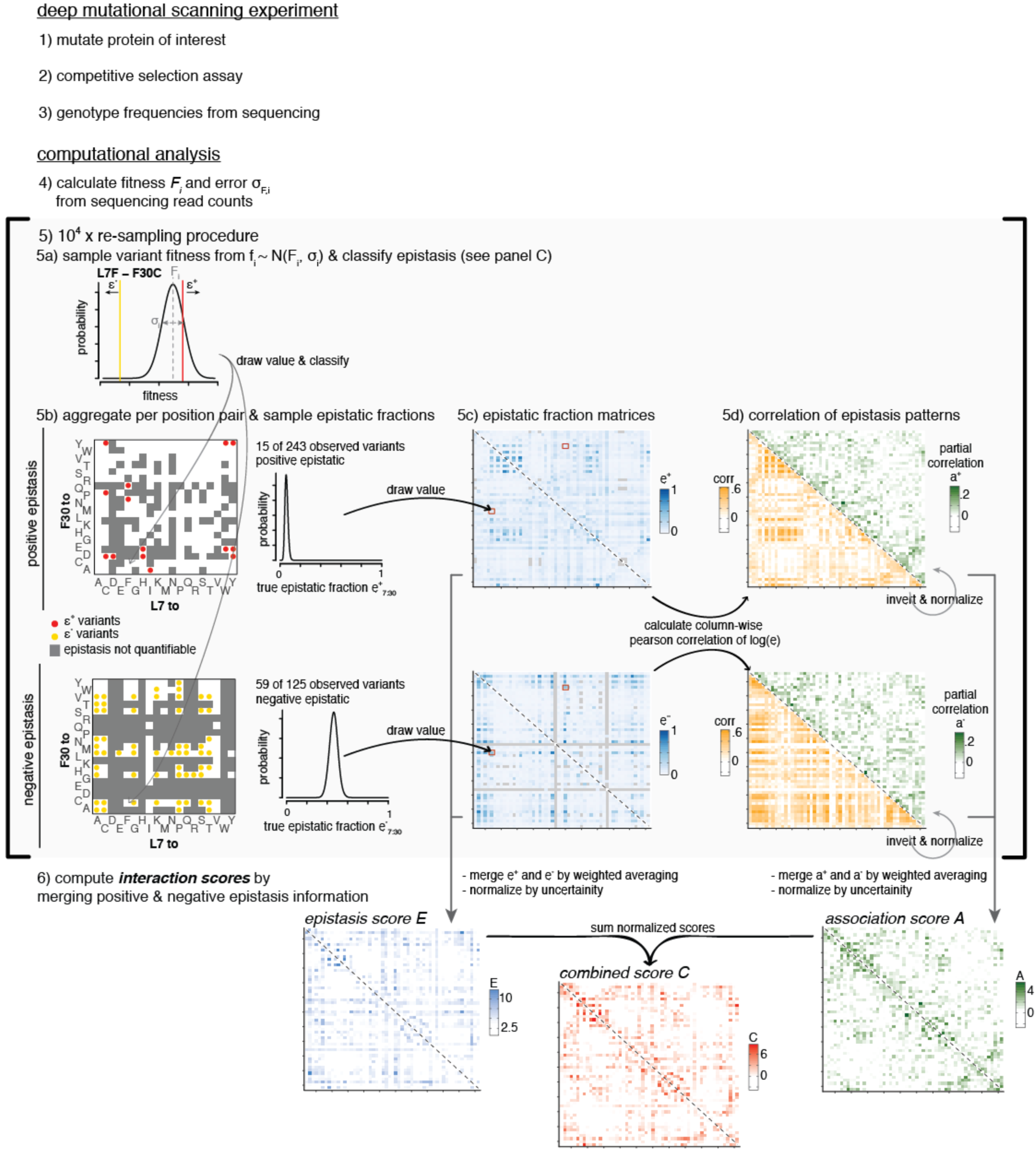
Deep mutational sequencing data to contact prediction workflow. Overview of workflow to predict interacting position pairs from deep mutational scanning datasets (see Methods and Results).

**Extended Data Figure 2:**
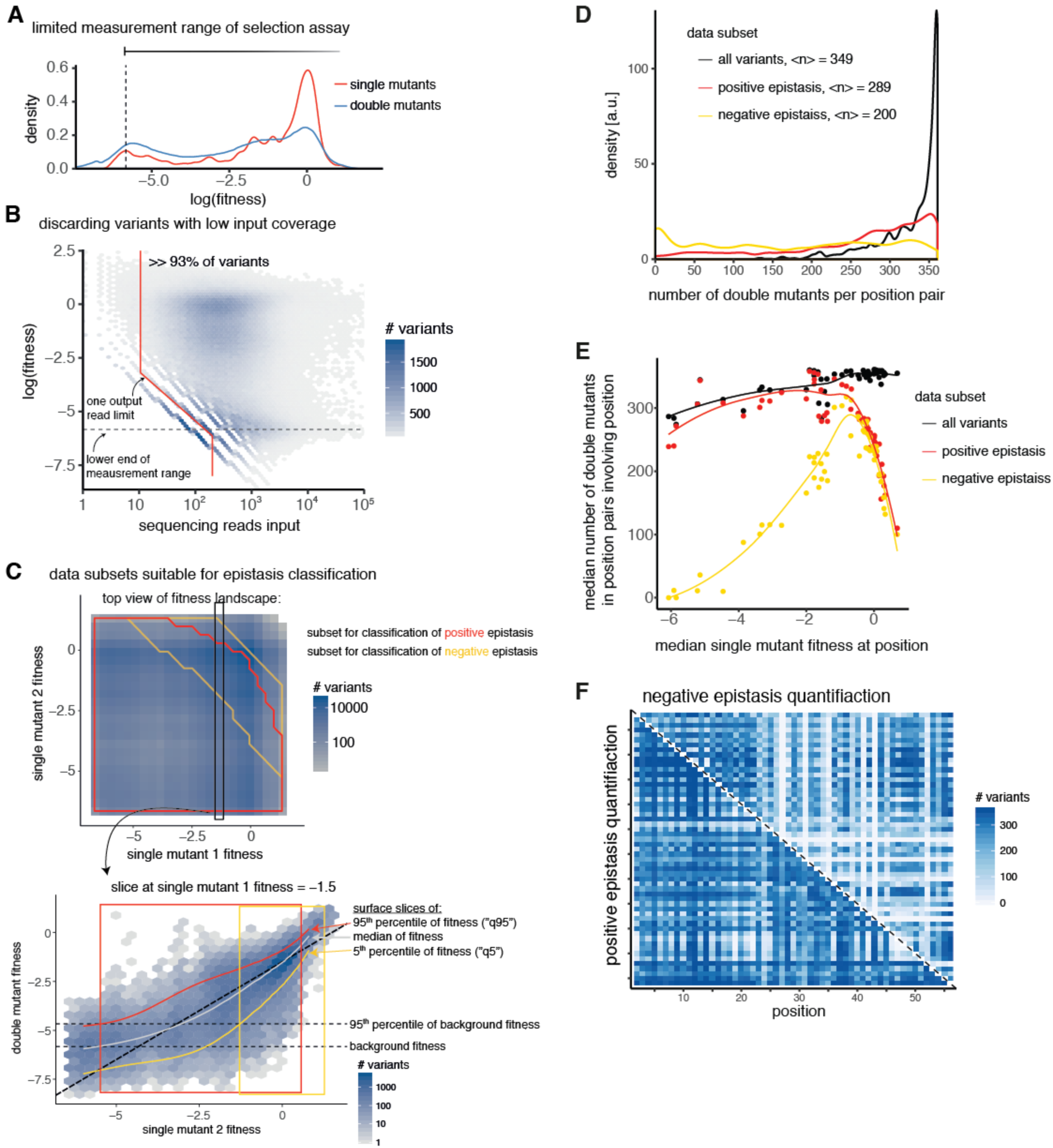
GB1 deep mutational scanning data processing. A. Distribution of fitness values for single and double mutant variants. Lower peak in distributions indicates lower limit of fitness assay measurement range (see Methods). B. Two-dimensional variant density showing dependency of fitness values on sequencing read counts in input library. For variants with very low coverage in the input library low fitness values cannot be accurately estimated. Red line shows sequence read cutoff used for variant inclusion (93% of variants included for downstream analysis). Horizontal dashed line indicates lower limit of fitness assay measurement range. C. Reliable quantification of positive and negative epistasis is limited to subsets of the data. Upper plot shows two-dimensional double mutant variant density in single mutant fitness space. Red outline shows single mutant fitness space that enables positive epistasis quantification. Yellow outline shows single mutant fitness space that enables negative epistasis quantification. Lower plot shows an example slice through fitness landscape at single mutant fitness = -1.5. Red, grey and yellow curves show slices through quantile fitness surfaces of 95^th^ percentile, median and 5^th^ percentile, respectively (see Figure 1B). Diagonal dashed line shows *double mutant fitness* = *single mutant* 2 *fitness* – 1.5 (expected fitness = observed fitness). Horizontal dashed lines give lower limit of fitness assay measurement range and the 95^th^ percentile of fitness values of variants dominated by background fitness effects. Red and yellow boxes indicate the range that includes 99% of variants within the slice that are suitable for positive or negative epistasis quantification, respectively. D. Distribution of number of double mutant variants across all position pair. Legend gives median number of double mutants per position pair for different data subsets. E. Relationship between median single mutant fitness at a position and the median number of double mutants observed in position pairs the position is involved in. Curves are loess smoothed. Across all variants, positions with stronger fitness effects show lower coverage of double mutants. Restrictions for quantification of positive epistasis additionally reduce coverage for positions with mostly neutral or positive effects. Finally, restrictions for quantification of negative epistasis strongly reduce coverage for positions with strong fitness effects, due to the lower measurement limit of the fitness assay. F. Number of double mutants for which positive (lower left triangle) or negative (upper right triangle) epistasis can be quantified per position pair plotted on the interaction matrix.

**Extended Data Figure 3:**
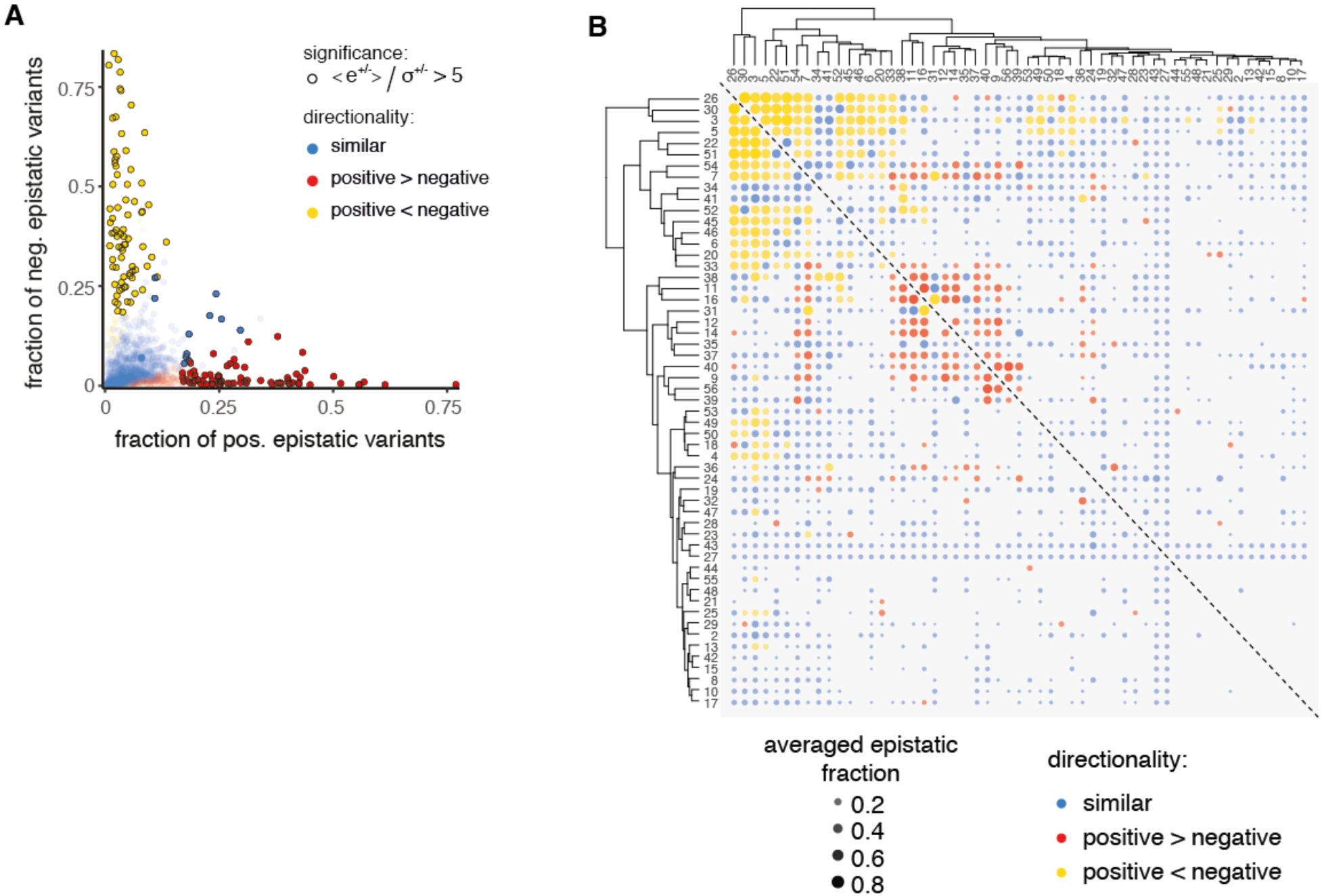
Positive and negative epistasis enrichments across position pairs. A. Position pair-wise fractions of positive and negative epistatic variants. Black circles mark position pairs with highly significant fractions (either positive or negative, or both); red dots: positive epistatic fraction significantly larger than negative epistatic fraction; yellow dots: negative epistatic fraction significantly larger than positive epistatic fraction; blue dots: no significant differences. B. Hierarchical cluster analysis of epistatic fraction patterns. Positions are clustered according to the Euclidean distance of their mean epistatic fractions (weighted average of positive and negative epistatic fractions, weights are inverse variances of fractions in resampling runs) to all other positions. Note that directionality of interactions (positive or negative fractions more significant) was not used for clustering but only marked post-analysis. Clustering shows two highly interconnected clusters of positions that interact mostly positively or negatively within each cluster but hardly any strong interactions are observed between the two clusters (with exception of positions 7 and 54).

**Extended Data Figure 4:**
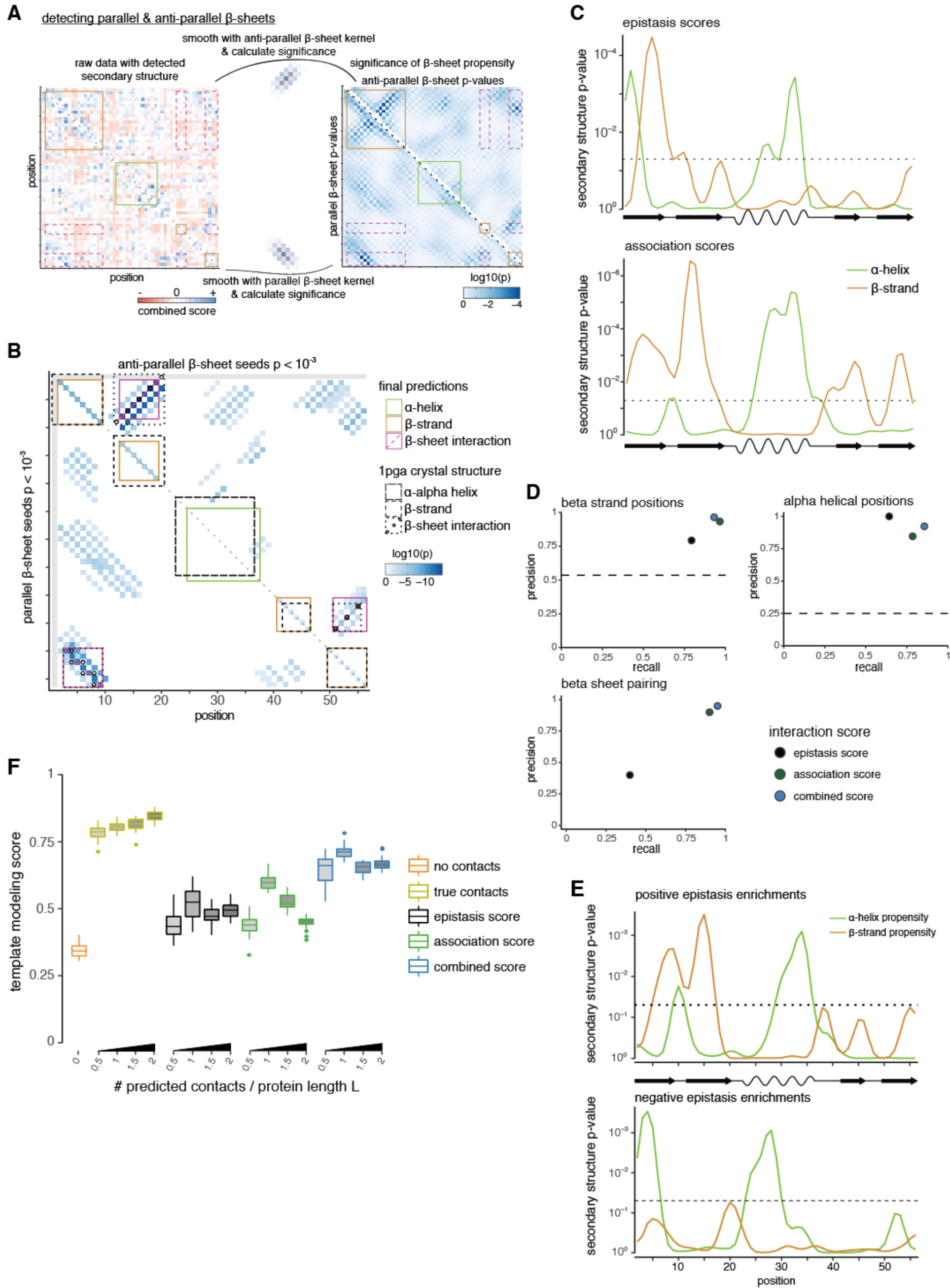
Secondary and tertiary structure prediction for GB1 domain. A. Detecting beta sheet pairing with two-dimensional kernel smoothing. Left plot shows raw combined score interaction matrix, with secondary structure element predictions (see Figure 4A,B) marked as squares along the diagonal (red – beta strand, green – alpha helix). Off-diagonal orange rectangles show potential regions of beta sheet pairing. Right plot: calculation of beta sheet pairing propensity with beta sheet kernels. Upper right triangle shows anti-parallel beta sheet propensity. Lower left shows parallel beta sheet propensity. B. Matrix of aggregated propensities of beta sheet pairing stretches (upper right – anti-parallel, lower left – parallel, p < 10^-3^) and the predictions for beta sheet pairing and secondary structure elements derived from them. In brief, predictions are performed by picking the highest propensity stretch that is consistent with predicted beta strands, if necessary modifying beta strand predictions (e.g. introducing an initially not predicted split between beta strands 1 and 2), then disregarding all stretches that conflict with the picked top-stretch. This procedure is repeated until no more beta sheet stretches with propensity P < 10^-3^ are left. Finally, beta strand predictions are updated such that only positions involved in a beta sheet interaction are retained. Reference elements from crystal structure are shown as comparison (lower triangle – parallel beta sheets, upper triangle – anti-parallel beta sheet, diagonal –secondary structure elements). C. Secondary structure propensity derived from kernel smoothing (red – beta strand, green – alpha helix) for *epistasis* (upper) and *association scores* (lower). P-values were derived by comparison to randomized datasets (see Methods). Dashed line indicates p = 0.05. D Precision and recall for beta strand, alpha helix and beta sheet predictions from *epistasis, association* and *combined scores* (in comparison to crystal structure). Dashed lines for beta strand and alpha helical positions give random expectation. Random expectation for beta sheet pairing precision is below 1%. E. Secondary structure propensities derived from local positive or negative epistatic enrichments. The upper panel shows secondary structure propensity derived from positive epistatic interactions, which are in line with secondary structure elements in the GB1 crystal structure (PDB entry 1pga). The lower panel shows secondary structure propensity derived from negative epistatic interactions, which are devoid of beta strand signals and instead show a three-helical pattern, which is reminiscent of the three-helical structure of the protein G A domain that binds albumin ^57^. F. Template modeling score of top 5% structural models compared to crystal structure 1pga and the dependency on number of predicted contacts used. “No contacts” – only restraints for secondary structure predicted by PSIPRED. “True contacts” – restraints derived from 0.5-2*L contacts (linear sequence separation greater than 5 positions, random subset), secondary structure elements and beta sheet interactions from crystal structure. All other: restraints derived from top 0.5-2*L contacts, secondary structure element and beta sheet interaction predictions from the three interaction scores, as indicated by color.

**Extended Data Figure 5:**
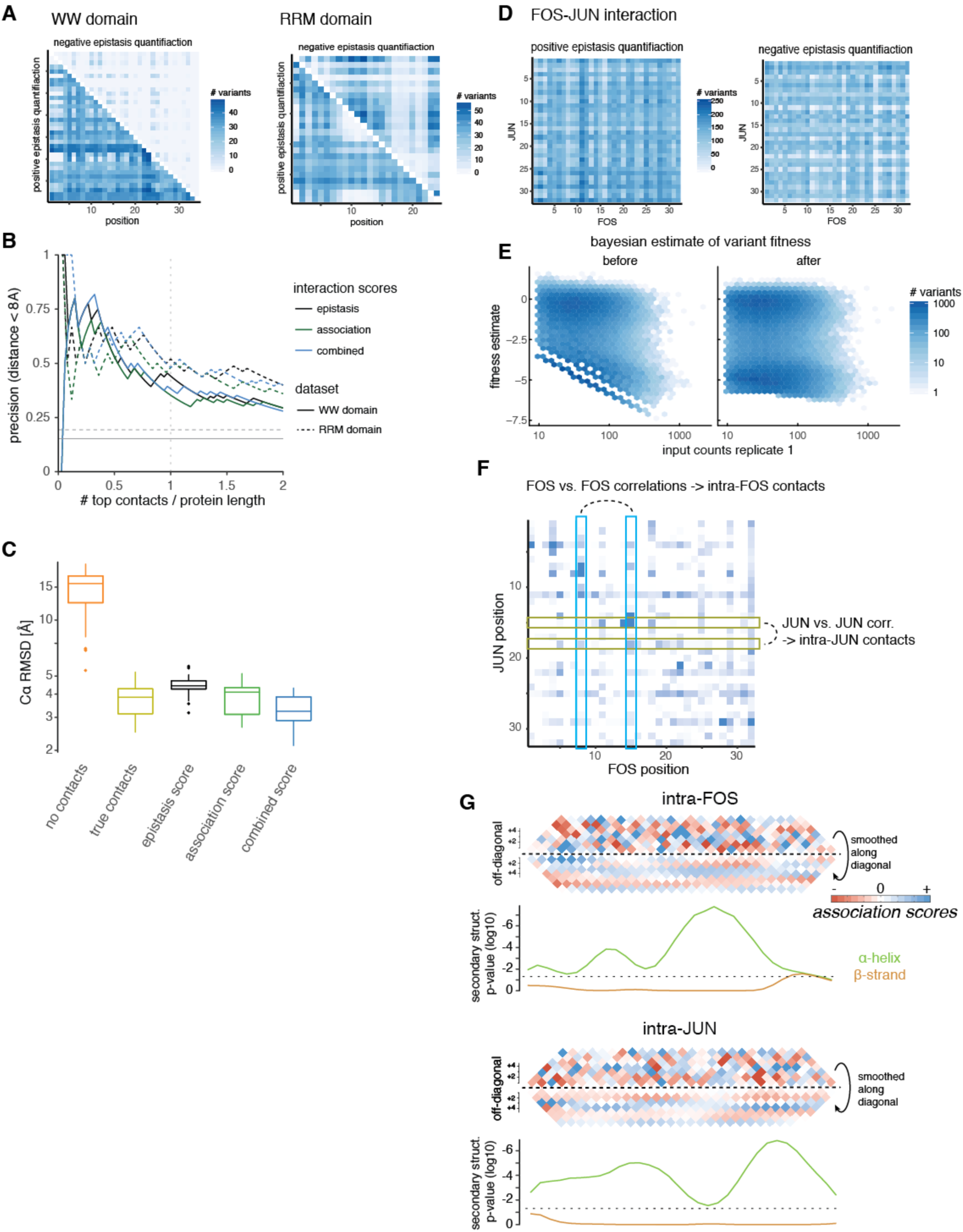
WW and RRM domain and FOS-JUN interaction dataset properties. A. Number of double mutants for which positive (lower left triangle) or negative (upper right triangle) epistasis can be quantified per position pair plotted on the interaction matrix for WW (left) and RRM (right) domains. B. Precision of interaction scores to predict direct contacts (distance < 8Å in reference structure) as a function of top scoring position pairs for WW and RRM domain interaction scores. Color denotes interaction scores, solid lines for WW domain, dashed lines for RRM domain. Grey horizontal lines give random expectation. Only position pairs with linear sequence separation greater than 5 amino acids are considered. C. Accuracy (*C*α root-mean-square deviation) of top 5% structural models of the WW domain (core positions 6-29) generated from deep mutational scanning data derived restraints compared to reference structure (PDB entry 1k9q). Structural models were generated in XPLOR-NIH by simulated annealing with restraints derived from top 17 tertiary contacts and secondary structure elements predicted by PSIPRED. No beta sheet pairing information was used. D. Number of double mutants for which positive (left) or negative (right) epistasis can be quantified per position pair plotted on the trans-interaction matrix of the FOS-JUN interaction. E. Bayesian estimation of fitness values in FOS-JUN interaction data. Mutants with low input sequencing coverage display limited measurement range and many dropouts (∽15% of variants without reads in output). Left panel shows original fitness distribution as function of input coverage in replicate 1, right panel shows Bayesian estimates of fitness as function of input coverage in replicate 1. F. Learning about intra-molecular contacts in FOS or JUN from epistatic pattern correlations. Column-wise correlation of epistatic patterns of the trans interaction score map serve to calculate intra-FOS *association scores* and thus reveal relationships between positions in FOS. Likewise, row-wise correlation of epistatic patterns reveal relationships between positions in JUN. G. Local interactions in intra-molecular *association scores* reveal secondary structures of protein interaction partners. Upper panels: Data above diagonal shows *association score* data close to the diagonal, i.e. local interactions. Data below the diagonal is smoothed with a Gaussian kernel to reveal interaction periodicity. Lower panels: Secondary structure propensities derived from kernel smoothing (see Figure 4A-C). Green indicates alpha helical propensity, orange indicates beta sheet propensity, p = 0.05 is indicated by dashed line.

**Extended Data Figure 6:**
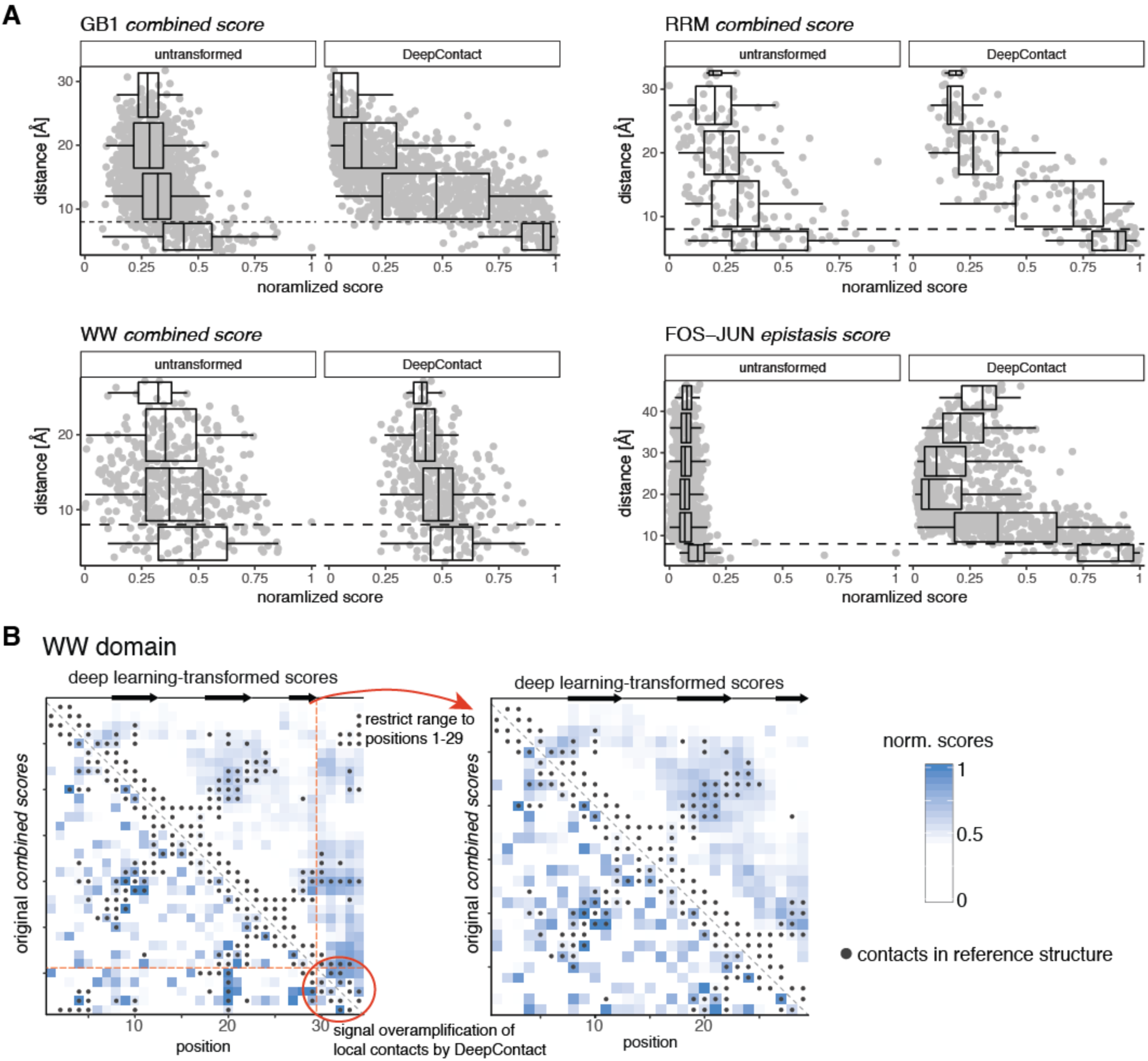
Deep learning improves contact prediction from deep mutagenesis data. A. Distance of position pairs as a function of interactions scores before (left panel, scores normalized to interval [0,1]) and after (right panel) transformation with *DeepContact* for the four datasets. Boxplots are spaced in distance intervals [0,8), [8,16), [16,24), [24,32), [32,40) and [40,48) Å. Dashed horizontal line indicates 8Å. B. Left: Full WW domain *combined score* interaction map before (lower left) and after (upper right) DeepContact transformation. DeepContact amplifies a signal from local contacts in the C-terminal region of the domain, thus concentrating the strongest transformed signal in this region. Removing positions 30-34 removes this artefact (right plot). Heat maps show interaction scores that have been normalized to have similar range. Grey dots show contacts (distance < 8Å) in reference structure.

**Extended Data Figure 7:**
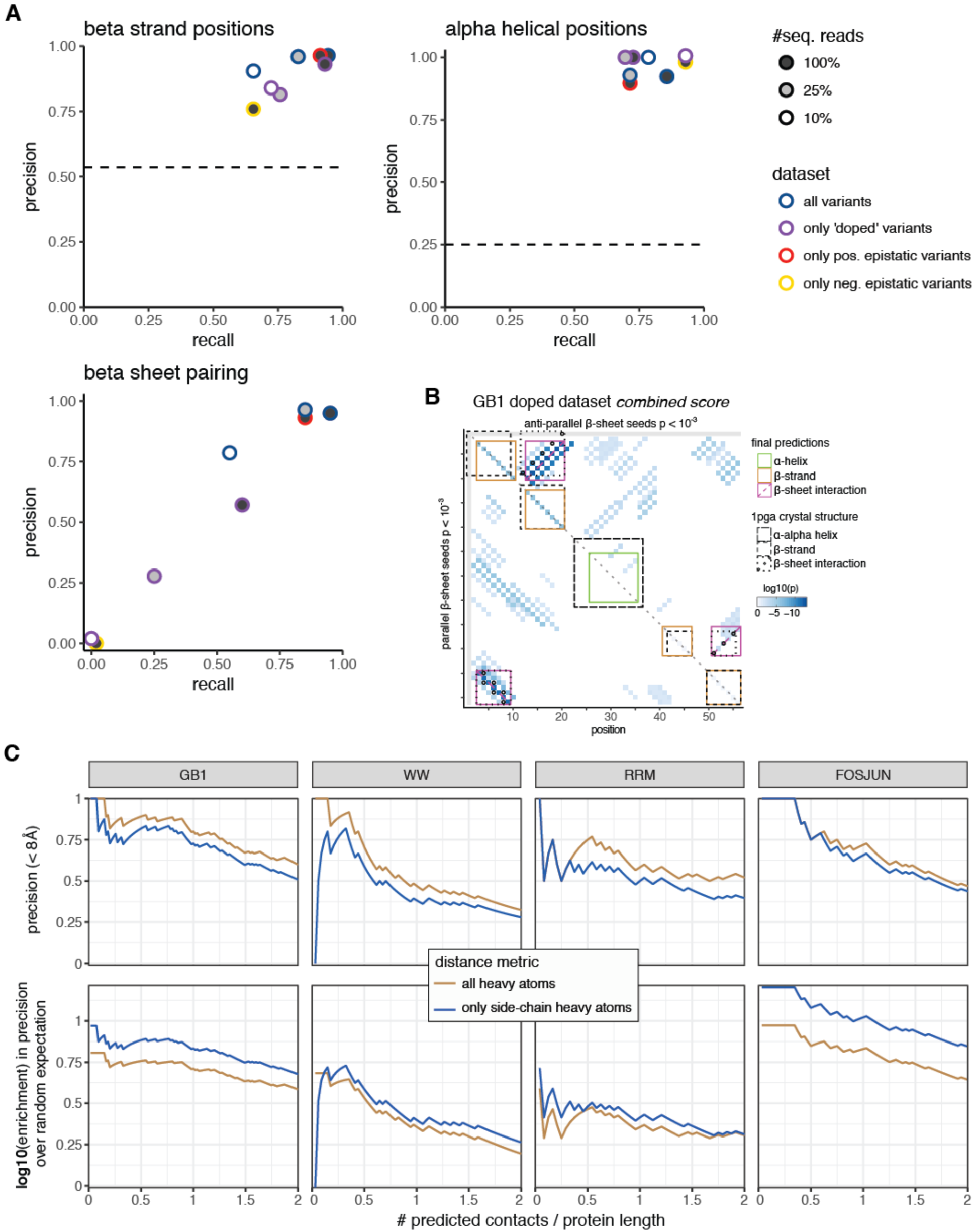
Distance metric comparison and secondary structure prediction for lower data-quality GB1 datasets. A. Precision and recall for beta strand, alpha helix and beta sheet predictions derived from combined scores of down-sampled GB1 datasets (in comparison to reference structure). Dashed lines for beta strand and alpha helical positions give random expectation. Random expectation for beta sheet pairing precision is below 1%. Note that some coinciding data points were slightly moved for better identifiability. B. Beta sheet pairing predictions for the doped GB1 dataset with 100% sequencing read coverage (cf. Extended Data Figure 4B). Beta sheet pairing between beta strands 1 and 2 is predicted in correct anti-parallel direction, but exact pairing of positions are off by 2; thus precision and recall of beta sheet pairing for doped GB1 dataset drops to ∽60% (see panel B). C. Differences in precision and enrichment over random expectation for all heavy atom or side-chain heavy atom distance metrics. As expected, using all heavy atoms (including backbone heavy atoms) increases precision of predicted contacts by about 10%. Restricting distance measurements to side-chain heavy atoms, however, increases precision over random expectation, often by more than 2-fold (note the log10-scale), indicating that side-chain distances are more informative for epistatic interactions. For these calculations, only position pairs with linear sequence separation greater than 5 amino acids were considered.

